# TDP-43 directly inhibits RNA accumulation in neurites through modulation of RNA stability

**DOI:** 10.1101/2025.02.25.640092

**Authors:** Charlie Moffatt, Ankita Arora, Katherine F. Vaeth, Bryan B. Guzman, Gurprit Bhardwaj, Audrey Hoelscher, Levi B. Gifford, Holger A. Russ, Daniel Dominguez, J. Matthew Taliaferro

**Affiliations:** Department of Biochemistry and Molecular Genetics, University of Colorado Anschutz Medical Campus, Aurora, CO, USA; Department of Pharmacology, University of North Carolina at Chapel Hill, USA; Diabetes Institute, University of Florida, USA; Department of Pharmacology and Therapeutics, University of Florida, USA; RNA Bioscience Initiative, University of Colorado Anschutz Medical Campus, Aurora, CO, USA; RNA Discovery Center, University of North Carolina at Chapel Hill, USA

## Abstract

The subcellular localization of hundreds of RNAs to neuronal projections allows neurons to efficiently and rapidly react to spatially restricted external cues. However, for the vast majority of these RNAs, the mechanisms that govern their localization are unknown. Here we demonstrate that the ALS-associated RNA binding protein TDP-43 primarily acts to keep RNAs out of neuronal projections. Using subcellular fractionation and single molecule RNA FISH we find that TDP-43 loss results in the increased neurite accumulation of hundreds of RNAs. These RNAs are highly enriched for known TDP-43 binding sites, suggesting that TDP-43 directly binds them to regulate their localization. We then identified precise regions within RNAs that mediate their TDP-43-dependent localization and interaction with TDP-43 using high-throughput functional assays in cells and high-throughput binding assays *in vitro*. We found that these regions also mediated TDP-43-dependent RNA instability, identifying the mechanism by which TDP-43 regulates RNA localization. ALS-associated mutations in TDP-43 resulted in similar RNA mislocalization phenotypes as did TDP-43 loss in human iPS-derived motor neurons. These findings establish TDP-43 as a direct negative regulator of RNA abundance in neurites and suggest that mislocalization of specific transcripts may occur in ALS patients.

## INTRODUCTION

Thousands of mRNAs are asymmetrically distributed in diverse cell types and across species. The correct localization of many of these mRNAs is critical to normal organism development and cellular function [1–3]. Localized transcripts enable local translation of proteins, which in turn allows rapid, localized responses to internal cues or external stimuli [4–8]. In particular, the elongated shape and large distances associated with neuronal morphology mean that RNA localization plays a key role in the ability of these cells to efficiently populate their subcellular proteomes to allow spatially restricted function [5,7].

The localization of these RNAs is regulated through the presence of features within the transcript, often called “zip codes” [9–11]. These cis-elements are recognized by trans-factors, often RBPs, that mediate transport [12–17]. Although thousands of RNAs are regulated in their spatial distributions, the detailed mechanism of the cis-elments and trans-factors involved in such control is known for only a handful [13,18]. For most localized RNAs, the processes that regulate their transport are unknown.

Mutations in RBPs that regulate RNA localization are associated with neurological disease [8,16,17,19–23]. Of these RBPs, TAR DNA binding protein 43 (TDP-43) has been subject of much work due to its association with amyotrophic lateral sclerosis (ALS) [19,20,24–27]. TDP-43 is perhaps best known as a regulator of alternative pre-mRNA splicing [28], and TDP-43-related defects in alternative splicing have been directly linked to ALS phenotypes [29]. TDP-43 also negatively regulates RNA stability, including of its own transcript[30], and defects in this process have also been linked to ALS phenotypes [31].

Multiple studies have provided evidence that TDP-43 also regulates RNA localization [32–36]. These studies have found that TDP-43 typically promotes RNA transport to neuronal projections. However, the RNAs that depend upon TDP-43 for proper localization in neurons remain largely unidentified. Furthermore, how TDP-43 recognizes these RNAs in order to mediate their localization similarly remains generally unknown.

Here, using subcellular fractionation, single molecule imaging, and multiple quantitative high-throughput binding, functional, and RNA metabolic assays, we find that direct interaction between TDP-43 and its RNA targets instead reduces the accumulation of those RNAs in neurites. We find that this is likely due to TDP-43 promoting the turnover of its RNA targets, preventing the neurite accumulation of these RNAs. We identify specific regions within these RNAs that are responsible for this effect as well as targeted mutations that abolish it. Finally, we find evidence of this effect in multiple primary and iPS-derived neuronal samples as well as evidence that TDP-43-mediated RNA localization is inhibited by ALS-associated mutations. Together, these findings establish new mechanisms by which TDP-43 regulates neuronal RNA metabolism.

## RESULTS

### Generation of TDP-43 knockout mouse neuronal cell lines

To quantify RNA localization in neuronal cell types, we and others have repeatedly used microporous transwell membranes [5,9–11,37–39]. The pores in these membranes are large enough to allow neurites to pass through, but cell bodies are restricted to the top of the membrane. After neurite growth, scraping the top of the membrane mechanically separates cells into cell body and neurite fractions (**Figure 1A**). By isolating RNA from each fraction and analyzing it using high-throughput sequencing, the relative abundance of an RNA in each fraction can be quantified.

**Figure 1.**
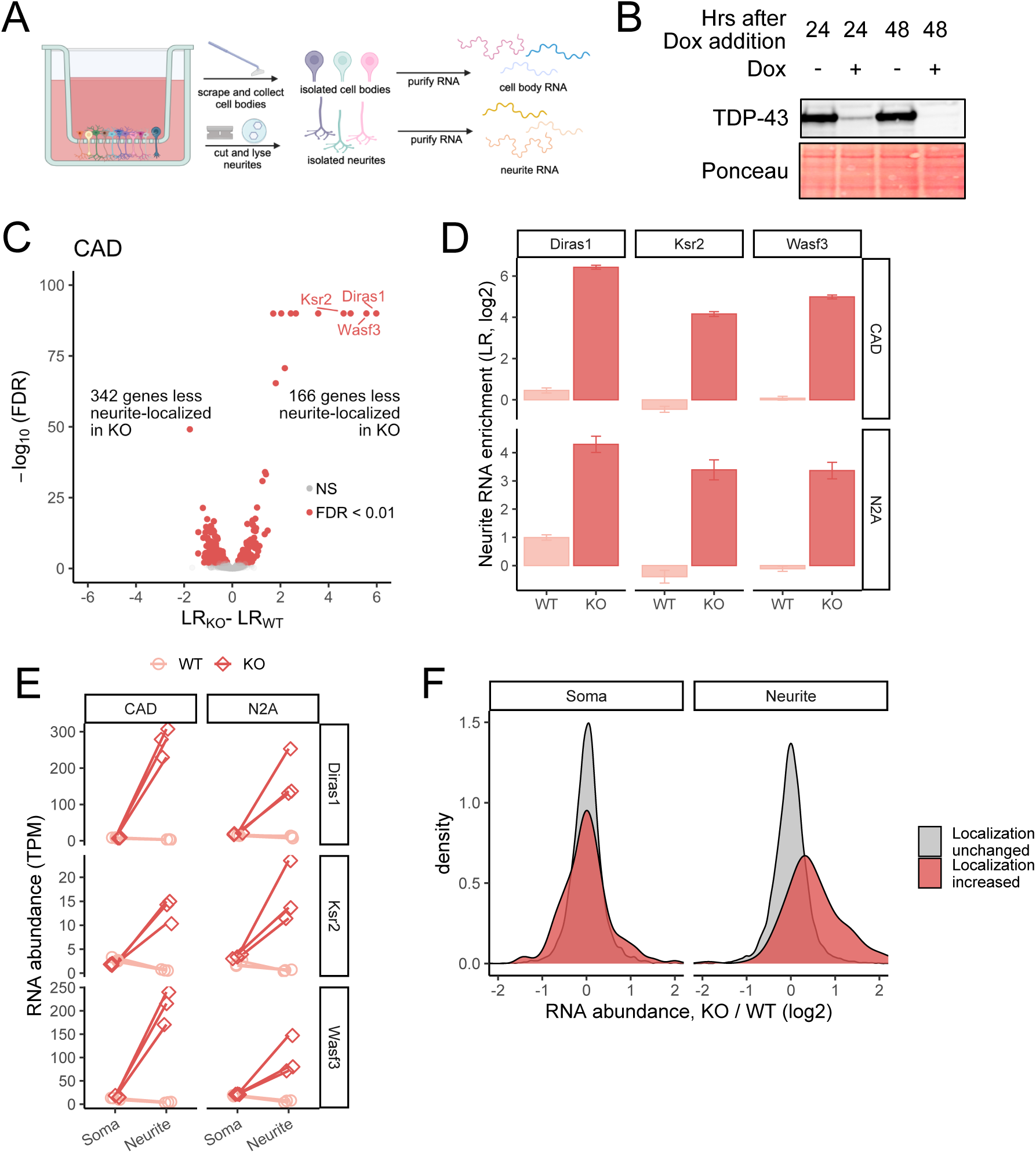
TDP-43 inhibits RNA localization to neurites. (A) Schematic of subcellular fractionation and RNA isolation setup. (B) Immunoblot of CAD conditional TDP-43 knockout cells demonstrating loss of TDP-43 expression following doxycycline addition. (C) Changes in RNA localization to neurites in CAD cells upon loss of TDP-43. (D) Changes in RNA localization to neurites of Diras1, Ksr2, and Wasf3 transcripts in CAD and N2A cells following TDP-43 loss. (E) Changes in abundances of Diras1, Ksr2, and Wasf3 transcripts in soma and neurite compartments following TDP-43 loss. (F) Changes in neurite and soma abundances in CAD cells for all transcripts following TDP-43 loss.

In order to identify RNAs whose localization to neurites depends on TDP-43, we first sought to create TDP-43 knockouts in two unrelated mouse neuronal cell lines, CAD and N2A, that have repeatedly been used to investigate RNA localization to neurites [9,17,37,38]. These cell lines contain a single loxP-flanked cassette in their genome, allowing efficient and controlled expression of transgenes [40].

Given that TDP-43 is essential for survival in many cell types [41], we reasoned that a simple knockout approach may not be viable. We therefore designed a genetic bypass approach whereby a single-copy TDP-43 transgene under the control of a doxycycline-off promoter was site-specifically integrated using cre/lox recombination. This HA-tagged transgene contained silent mutations that rendered it immune to guide RNAs that were then used to inactivate endogenous TDP-43 alleles with CRISPR/Cas9. Using immunoblotting, we then screened for clones in which endogenous TDP-43 expression was ablated (**Figure S1A**). In these clones, the only functional TDP-43 allele remaining was the transgene, which could then be inactivated with the addition of doxycycline. After 48 hours of doxycycline treatment, TDP-43 was undetectable by immunoblotting (**Figure 1B**). Extended incubation in doxycycline for 2 weeks resulted in cell death, likely pointing again to TDP-43 being essential. All further experiments were done following 48 hours of doxycycline treatment. Following acute TDP-43 depletion at the 48 hour timepoint, we observed that neurites were slightly shorter (**Figure S1B**), but did not observe noticeable cell death.

### Hundreds of RNAs are mislocalized in TDP-43 knockout cells

To identify RNAs that were mislocalized following TDP-43 loss, we fractionated these cells into cell body and neurite fractions following 48 hours of doxycycline treatment (knockout) or the omission of doxycycline (wildtype). For each RNA, we then calculated a Localization Ratio (LR), which we defined as the log2 of its relative abundance in the neurite fraction compared to its relative abundance in the cell body fraction. Higher (positive) LR values therefore indicate greater enrichments of an RNA in neurites while lower (negative) values indicate depletion from neurites. To maximize the generality of these experiments and reduce cell line-associated idiosyncracies, we performed them both in CAD and N2A cells.

We first assayed the efficiency of the cell body / neurite fractionation by calculating LR values for RNAs known to be enriched in neurites. These include RNAs encoding ribosomal proteins and components of the electron transport chain [37]. Reassuringly, we found that these RNAs were consistently neurite-enriched in our data, indicating its reliability (**Figure S1C**). We further assayed the quality of the data using PCA and hierarchical clustering. We found that replicates clustered tightly with each other and that samples were cleanly separated first by subcellular location (cell body vs. neurite) and then by TDP-43 status (wildtype vs. knockout) (**Figure S1D-G**).

We then moved to the identification of RNAs whose LR value significantly differed between wildtype and knockout samples. In both cell lines, we identified hundreds of RNAs that were either more or less neurite-enriched in knockout cells compared to wildtype controls (**Figure 1C, Figure S1H, Table S1, Table S2**). When we compared results from the two cell lines, we found the same RNAs were similarly mislocalized in both, again giving us confidence in the results (**Figure S1I, J**).

Interestingly, we found that a handful of RNAs, exemplified by Diras1, Ksr2, and Wasf3, were 20-30 fold more neurite-enriched in knockout cells compared to wildtype cells, indicating that TDP-43 acts to keep them out of neurites (**Figure 1C, D**). We confirmed this using qRT-PCR (**Figure S1K**). Because our LR metric is a ratio, in principle, the increased LR value in knockout cells could be due to either an increase in neurite abundance or a decrease in soma abundance. To distinguish between these possibilities, we compared expression values from each subcellular fraction. The abundance of these RNAs in soma samples were approximately equal in wildtype and knockout samples. In contrast, neurite abundances were 10-30 fold higher in knockout samples (**Figure 1E**). In fact, this trend held throughout the hundreds of RNAs that displayed increased neurite enrichments in knockout cells (**Figure 1F**). From these results, we conclude that the increased neurite enrichments of these RNAs observed in knockout cells is primarily due to a neurite-specific increase in their abundance.

### Validation of changes in subcellular RNA localization using single molecule RNA FISH

Given that we observed similar results in TDP-43-dependent RNA localization changes in both CAD and N2A cell lines (**Figure 1C, S1H-J**), for simplicity, we chose to focus on the CAD cell line for further experiments. To independent validate changes in RNA localization observed via RNAseq, we measured the localization of Ksr2 RNA using single molecule fluorescence in situ hybridization (smFISH) [42]. We counted transcript abundances in soma and neurites in wildtype and knockout cells. We found that Ksr2 became significantly more neurite-enriched in TDP-43 knockout cells, consistent with the subcellular fractionation and RNAseq results (**Figure 2A, B**).

**Figure 2.**
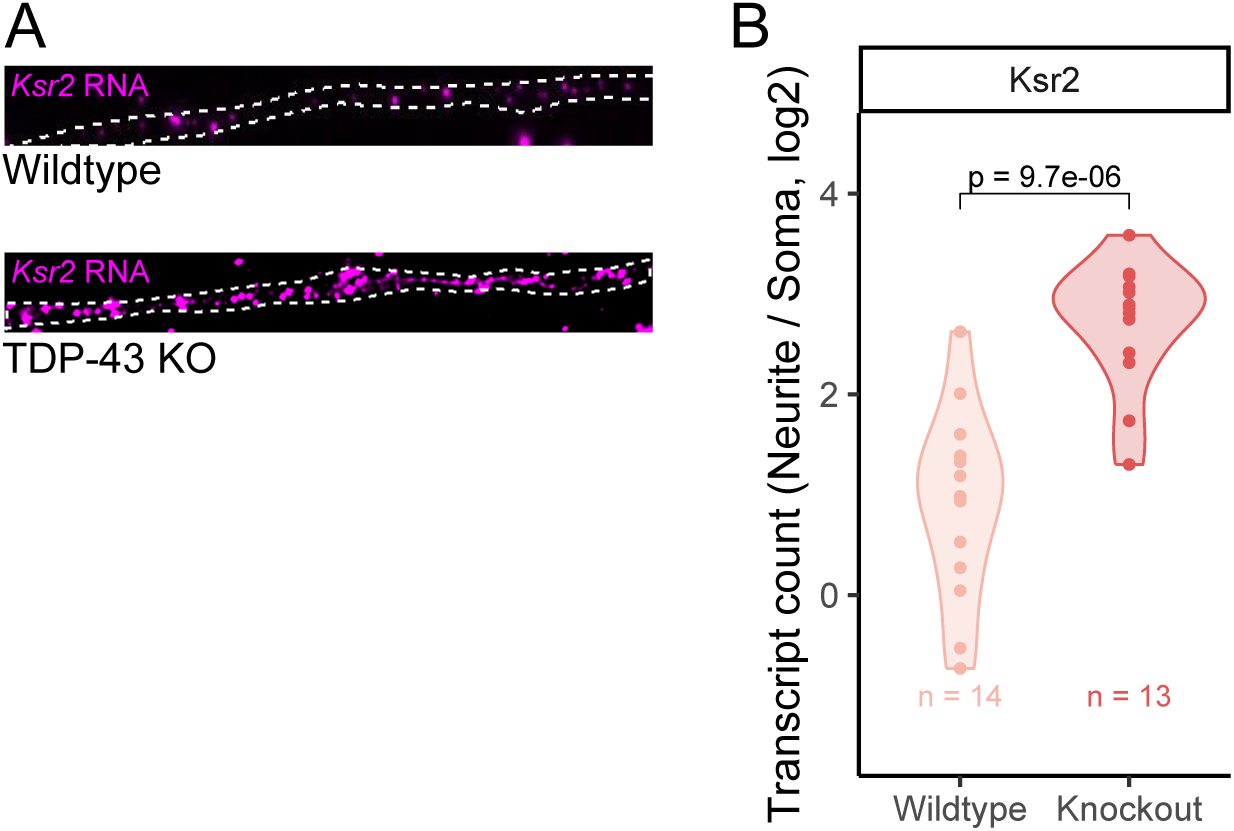
Validation of RNA mislocalization in TDP-43 knockout cells using smFISH. (A) Representative images of CAD neurites from wildtype and TDP-43 knockout cells. Ksr2 RNA is visualized using smFISH (magenta). (B) Quantification of smFISH results. P values were calculated using a Wilcoxon ranksum test.

### TDP-43 RNA localization targets are highly enriched in RNA sequences directly bound by TDP-43

The preferred binding motif of TDP-43 has been extensively studied through a variety of methods including CLIP-seq and RNA Bind-N-Seq [28,43], and has been repeatedly identified as a repeat of a GU dinucleotide with an occasional tolerance for adenosine between two uracil residues (**Figure 3A**). If the observed TDP-43-dependent localization of specific transcripts was due to TDP-43 directly binding them, we would expect these transcripts to contain enrichments of these RNA motifs.

**Figure 3.**
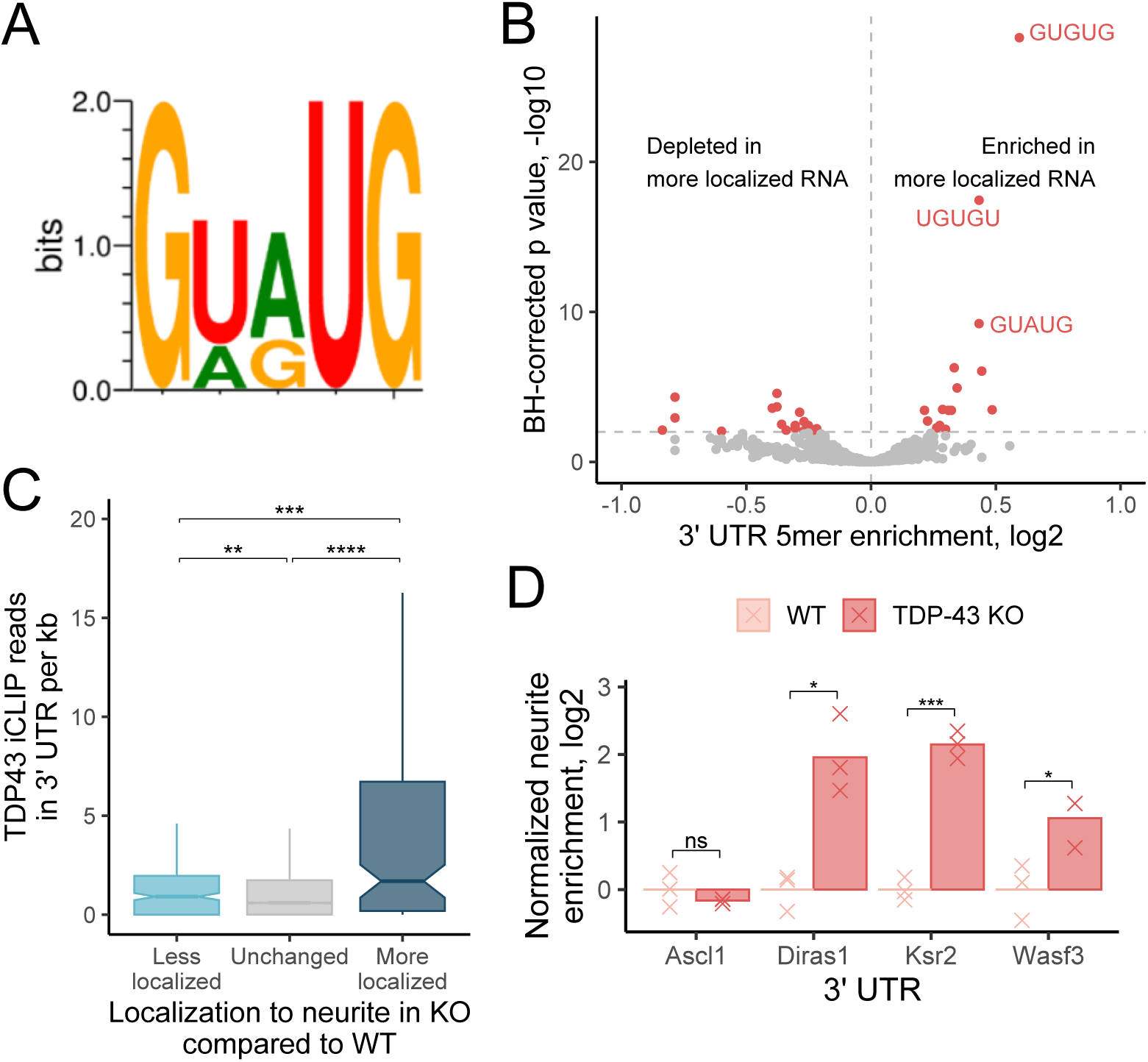
TDP-43 directly regulates its RNA localization targets. (A) Preferred RNA motif for TDP-43 binding. (B) Enrichment of all 5mers in the 3′ UTRs of RNAs that show increased neurite-enrichment following loss of TDP-43. Those that are known to be bound by TDP-43 are explicitly marked. (C) Distribution of TDP-43 CLIP-seq reads in the 3′ UTRs of RNAs that became less neurite-enriched, RNAs whose localization did not change, and those that became more neurite-enriched in TDP-43 knockout cells. (D) Differences in neurite localization for a reporter transcript containing the indicated 3′ UTRs in wildtype and TDP-43 knockout cells. In panel C, P-values were calculated using a Wilcoxon ranksum test. In panel D, P-values were calculated using a t test. NS (not significant) represents p >0.05, * p < 0.05, ** p < 0.01, *** p < 0.001, and **** represents p < 0.0001.

To test this, we compared the 3′ UTR sequences of the RNAs that displayed increased neurite localization in knockout cells to the 3′ UTR sequences of RNAs whose localization was not TDP-43-sensitive. We found that the top three 5mers enriched in the RNAs that displayed increased neurite localization were perfect matches for 5mers known to be bound by TDP-43 (GUGUG, UGUGU, GUAUG) (**Figure 3B**). These 5mers were not enriched in the 3′ UTRs of RNAs that displayed decreased neurite localization in knockout cells (**Figure S2A**). These results suggest that the RNAs that become more neurite-enriched upon TDP-43 loss are directly bound by TDP-43 in their 3′ UTRs. Conversely, those that become less neurite-enriched are not directly bound by TDP-43, and therefore their mislocalization is likely not directly due to a loss of TDP-43 binding.

The enrichments of the TDP-43 binding motifs in the more-localized RNAs were specific to their 3′ UTRs as they were not found in their coding sequences or 5′ UTRs (**Figure S2B, S2C**). However, the motifs were strongly enriched in the 3′ UTRs of the human orthologs of the more-localized RNAs (**Figure S2D**), and the 3′ UTRs of the more-localized RNAs were significantly more conserved than expected (**Figure S2E**). These results suggest that their regulation by TDP-43 may be conserved.

To look for more direct evidence of TDP-43 binding these RNAs, we analyzed a TDP-43 CLIP-seq dataset derived from mouse brain tissue [28]. We found that the 3′ UTRs of RNAs that became more neurite-enriched in knockout cells were approximately 3.5 times more likely to contain a TDP-43 CLIP-seq peak and contained a 3.5 fold higher density of TDP-43 CLIP-seq reads compared to TDP-43-insensitive RNAs (**Figure 3C, S2F**). The direct binding of the 3′ UTRs of these RNAs by TDP-43 was again conserved as the 3′ UTRs of their human orthologs were also highly enriched for direct binding in a human TDP-43 CLIP-seq experiment (**Figure S2G**) [44,45].

RNAs that became less-neurite enriched in knockout cells were again significantly less associated with TDP-43 binding in these CLIP-seq datasets (**Figure 3C, S3F**). Combined with the motif enrichment findings, these results suggest that the primary direct role for TDP-43 in the regulation of RNA localization in neuronal cells is to keep RNAs out of neurites. They further suggest that TDP-43 accomplishes this through binding specific sequences in the 3′ UTRs of its target RNAs.

### 3′ UTRs of TDP-43 RNA localization targets are sufficient to drive TDP-43-sensitive RNA localization

To directly experimentally test whether TDP-43-dependent changes in RNA localization were dependent upon the 3′ UTRs of the mislocalized transcripts, we turned to a reporter assay. In this experiment, the ratio of two reporter RNAs is measured in soma and neurite RNA samples using qRT-PCR. Comparing these ratios across the subcellular locations allows the quantification of the relative neurite-enrichment of the two reporters. By asking how that relative enrichment changes depending on the 3′ UTR appended onto one of the reporters, we can quantify the effect of that 3′ UTR on RNA localization (**Figure S2H**).

We appended the 3′ UTRs from three TDP-43-sensitive RNAs, Diras1, Ksr2, and Wasf3, onto our reporter. As a control, we also fused the 3′ UTR from a TDP-43-insensitive RNA, Ascl1. We then measured the neurite-enrichment of these reporters in wildtype and knockout cells. The localization of the control Ascl1 reporter was unaffected by TDP-43 loss. However, the other three reporters become strongly more neurite-enriched following TDP-43 loss, indicating that these 3′ UTRs contain elements that regulate RNA localization in a TDP-43-dependent manner and that TDP-43 acts to keep them out of neurites (**Figure 3D**).

### TDP-43 regulates ribosomal protein mRNA transport, but likely indirectly

Previous studies found that ribosomal protein mRNAs were aberrantly depleted from neurites in both TDP-43 knockdown primary mouse cortical neurons as well as in samples from ALS patients [36]. We also found that ribosomal proteins were significantly aberrantly depleted from neurites in TDP-43 knockout cells (**Figure S2I**), demonstrating that our CAD inducible knockout system is faithfully reporting on phenomena happening in primary cells. However, as a class, ribosomal protein mRNAs are significantly depleted for TDP-43 CLIP-seq peaks, suggesting that this effect may not be due to a loss of direct binding between TDP-43 and these mRNAs (**Figure S2J**).

### Design of massively parallel assay to identify sequence elements that regulate RNA localization in a TDP-43-dependent manner

Given that we identified full length 3′ UTRs that regulate RNA localization in a TDP-43-dependent manner, we wanted to know what sequences within these 3′ UTRs were responsible for this effect. To do this, we used a massively parallel reporter assay (MPRA) to simultaneously test the ability of thousands of discrete RNA sequences drawn from endogenous 3′ UTRs to mediate TDP-43-dependent RNA localization (**Figure 4A, Supplemental files 1-2**). We selected 15 3′ UTRs for this experiment. Of these, 9 came from RNAs that were more localized in TDP-43 knockout cells, 4 came from RNAs that were less localized in knockout cells, and 2 came from RNAs whose localization was not reliably affected by TDP-43 loss (**Figure S3A**).

**Figure 4.**
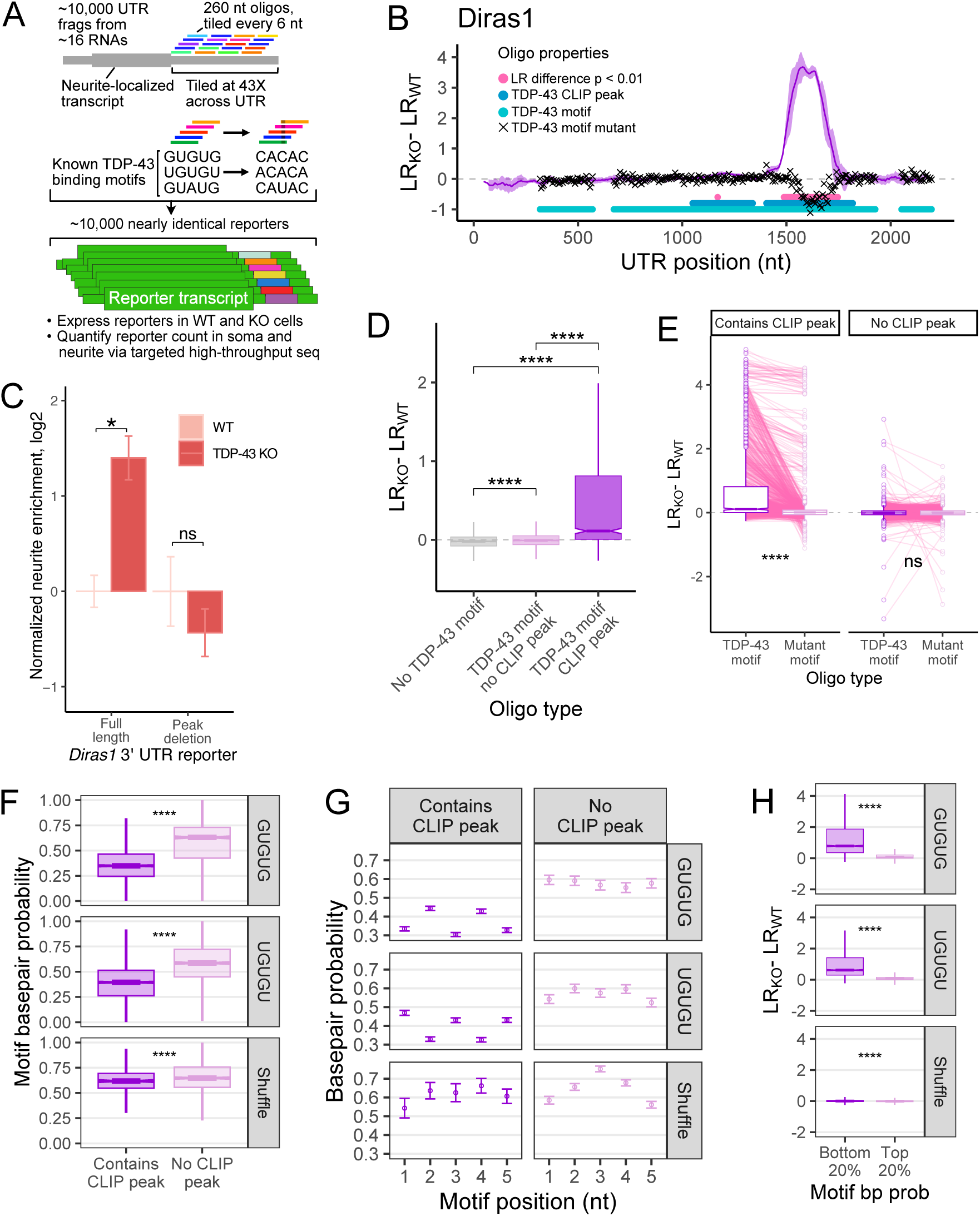
TDP-43 regulates RNA localization through the recognition of discrete sequence motifs in 3′ UTRs. (A) Design of MPRA for the interrogation of TDP-43-mediated RNA localization. (B) Differences in neurite enrichment between wildtype and knockout cells for all oligos that tile across the Diras1 3′ UTR. Oligos that contain a TDP-43 motif are represented by a light blue circle, those that contain a TDP-43 CLIP-seq peak are represented by a dark blue circle, and oligos with mutated TDP-43 motifs are represented by a black x. (C) Difference in neurite-enrichment in wildtype and knockout cells for reporters that contain the entire Diras1 3′ UTR (left) or the entire 3′ UTR lacking the peak of TDP-43 dependency identified in B (right). For panel C, P values were calculated using a t test. (D) Differences in RNA localization to neurites between wildtype and knockout cells for reporters containing oligos that do not have a TDP-43 motif (left), contain a motif but do not lie within a CLIP-seq peak (middle), or contain a motif and lie within a CLIP-seq peak (right). (E) Changes in neurite-localization between wildtype and knockout cells for reporter RNAs with oligos that contain TDP-43 motifs and their companion oligos in which those motifs were mutated. (F) Basepair probabilities for motifs in oligos that lie within or outside of TDP-43 CLIP-seq peaks. (G) As in F, but with basepair probabilities for individual nucleotides shown. (H) Differences in neurite RNA localization between wildtype and knockout cells for reporters with oligos that contain TDP-43 motifs that were highly single stranded (left) or highly double stranded (right). For panels D-H, P values were calculated using Wilcoxon ranksum tests. NS (not significant) represents p >0.05, * p < 0.05, ** p < 0.01, *** p < 0.001, and **** represents p < 0.0001.

We designed and synthesized approximately 10,000 different 260nt oligonucleotides (oligos) that tiled across each 3′ UTR, with 6nt of spacing between neighboring oligos (**Figure 4A**). In addition to these oligos, for any oligo that contained a TDP-43 motif (defined as UGUGU, GUGUG, or GUAUG), a companion oligo was included in which that motif was mutated (to ACACA, CACAC, and CAUAC, respectively). These oligos were integrated into the 3′ UTR of a reporter transcript in a plasmid, to create approximately 10,000 different reporters, each of which is identical except for the identity of the inserted oligo [9]. These plasmids were then transfected into wildtype and knockout cells, and the localization of the reporters was assayed via subcellular fractionation and UMI-aware RNA amplicon sequencing targeted against the reporter-embedded oligos. The abundance of each oligo was therefore quantified in the neurites and soma of wildtype and knockout cells (**Table S3**).

To ensure the quality of MPRA data, we performed PCA and hierarchical clustering analysis on the oligo abundance counts from each compartment and genotype. Oligo abundances clustered by subcellular compartment then genotype, indicating the data was of high quality (**Figures S3B, C**). We then quantified the neurite enrichment (LR) of each reporter-embedded oligo by comparing its normalized counts in the neurite RNA to its normalized counts in the soma as before.

Comparing LR values for each oligo across genotypes therefore allowed us to identify discrete 260nt RNA sequences that were sufficient to make the localization of an RNA TDP-43-sensitive.

### Characteristics of sequence elements that regulate RNA localization via TDP-43

We then examined the patterns of change in LR between wildtype and knockout cells on the gene level. To visualize the localization activity of the oligos derived from specific regions of each 3′ UTR, we plotted the rolling mean of the difference in LR by the position of each oligo in the 3′ UTR. Focusing on Diras1, oligos derived from most regions of the 3′ UTR do not exhibit any change in LR. However, oligos from one region made the localization of reporter transcripts highly sensitive to TDP-43 loss (**Figure 4B**). Interestingly, this region lined up perfectly with a previously known site of interaction with TDP-43 identified via CLIP-seq in mouse brain [28]. Similar results were seen with 3′ UTRs from other TDP-43-sensitive RNAs (**Figure S3D-E**), but no peaks of activity were observed in 3′ UTRs of TDP-43-insensitive RNAs (**Figure S3F**).

When the Diras1 3′ UTR region containing this peak of activity was deleted from the full length 3′ UTR, the localization of the reporter RNA was no longer TDP-43-sensitive, indicating that this 260nt region is both necessary and sufficient for regulation by TDP-43 (**Figure 4C**). Furthermore, oligos that made the reporter become more neurite-enriched were significantly more conserved than expected, suggesting that the ability of TDP-43 to regulate localization of their parent RNAs may also be conserved (**Figure S3G**). These results suggest that TDP-43 directly interacts with these 3′ UTR regions to regulate RNA localization.

To expand this analysis more generally, we separated oligos based on whether or not they contained a TDP-43 sequence motif and whether or not they contained a known site of TDP-43 interaction as identified by CLIP-seq [28]. Although many oligos contained at least one TDP-43 sequence motif, only a small subset of these were observed to actually be bound by TDP-43 by CLIP-seq (**Figure 4B**). While the presence of a TDP-43 motif alone was associated with increased neurite abundance in knockout cells, the size of this effect was very modest (**Figure 4D**). However, if that oligo also lay within a TDP-43 CLIP-seq peak (i.e. the motif was occupied by TDP-43 *in vivo*), the effect size was much larger. These results suggest the factors that determine whether or not a TDP-43 motif will be occupied by the protein are contained within the local (∼260nt) sequence context and recapitulated in the MPRA.

### The ability of TDP-43 to regulate RNA localization requires a discrete, canonical sequence motif

Although TDP-43 motifs (UGUGU, GUGUG, and GUAUG) were enriched in the 3′ UTRs of affected RNAs, this does not necessarily mean they are required for TDP-43 function upon those RNAs. To directly test the necessity of these motifs, we compared the change in LR values for wildtype, motif-containing oligos to their motif-mutated counterparts (**Figure 4A**). We found that mutation of TDP-43 motifs within TDP-43-sensitive oligos drastically reduced the ability of the oligo to regulate RNA localization in a TDP-43-dependent manner (**Figure 4A, 4E, S3D-F, H**). However, this effect was dependent upon the motif lying within a TDP-43 CLIP-seq peak (i.e. being occupied by the protein *in vivo*) (**Figure 4E, S3I**). These results further indicate that sequence motifs themselves are not alone sufficient and that additional contextual features in the surrounding RNA sequence distinguish functional and non-functional motifs.

### TDP-43’s occupancy and functional impact on an RNA is determined by RNA secondary structure

We wanted to further investigate what distinguished occupied (i.e. functional) from unoccupied (i.e. nonfunctional) TDP-43 motifs in these UTRs. Given that the distinguishing factor must be contained within the local RNA context due to the design of the MPRA, we reasoned that RNA secondary structure may be involved. Many RNA binding proteins have a preference for single stranded RNA [43], and we reasoned that TDP-43 may more efficiently recognize its sequence motif when the motif is single stranded [46].

To test this, we computationally folded each 260nt RNA sequence used in the MPRA (see Methods) [47]. We then binned oligos depending on whether or not they contained a TDP-43 CLIP-seq peak and calculated the probability that TDP-43 motifs within those oligos were participating in basepairing interactions. We found that TDP-43 motifs from oligos lying within CLIP-seq peaks were significantly more single stranded than motifs from oligos that were not within CLIP-seq peaks (**Figure 4F**). Furthermore, this effect was specific to TDP-43 motifs as it was clearly dampened for shuffled, control motifs (**Figure 4F**). We then analyzed the single-stranded character of each nucleotide of the motif individually. We found that guanosine nucleotides within the motif showed a particular preference for being unpaired when the motif was occupied (**Figure 4G**).

If RNA secondary structure were a driving factor in determining whether or not a TDP-43 motif was able to regulate RNA localization, we should be able to predict TDP-43-dependent changes in RNA localization based solely on the basepairing properties of the motif. To test this, we ranked all 260nt oligos in the MPRA by the basepairing probability of the TDP-43 motifs they contained. We then compared the LR values of the top and bottom 20% of oligos ranked by motif basepair probability. We found that oligos with single stranded motifs were much more likely to drive TDP-43-dependent changes in RNA localization than those with motifs that were more paired. Again, this effect was dependent upon the motif itself as control, scrambled motifs had much smaller effects (**Figure 4H**).

### Sequences that regulate RNA localization via TDP-43 in cells are directly bound by TDP-43 in vitro

The relationship between TDP-43 CLIP-seq peaks and oligos with TDP-43-sensitive LR values suggested that direct TDP-43 binding may be critical in the localization of these RNAs. To directly assay the affinity of TDP-43 for the same pool of RNA sequences used in the MPRA, we used RNA Bind-n-seq (RBNS) with purified recombinant TDP-43 [43,48]. By comparing results from the MPRA and RBNS, we are able to draw direct connections between TDP-43-dependent RNA localization activity and TDP-43 binding for the same defined pool of thousands of discrete 260mer RNA sequences.

With RBNS, the relative *in vitro* affinity (R) of a protein for any RNA within a complex pool can be calculated by comparing the abundances of each RNA in input and protein-bound samples (**Figure 5A, Table S4**). In these experiments, R is defined as the relative frequency of a sequence in the protein-bound sample divided by its relative frequency in the input sample. Reassuringly, we found that R values for all oligos calculated using different concentrations of TDP-43 clustered together, demonstrating the reproducibility of the RBNS assay (**Figure S4A**). Furthermore, as expected, we found that oligos with increasing numbers of TDP-43 motifs displayed higher affinities for TDP-43 (**Figure S4B**). These results gave us confidence in the RBNS data.

**Figure 5.**
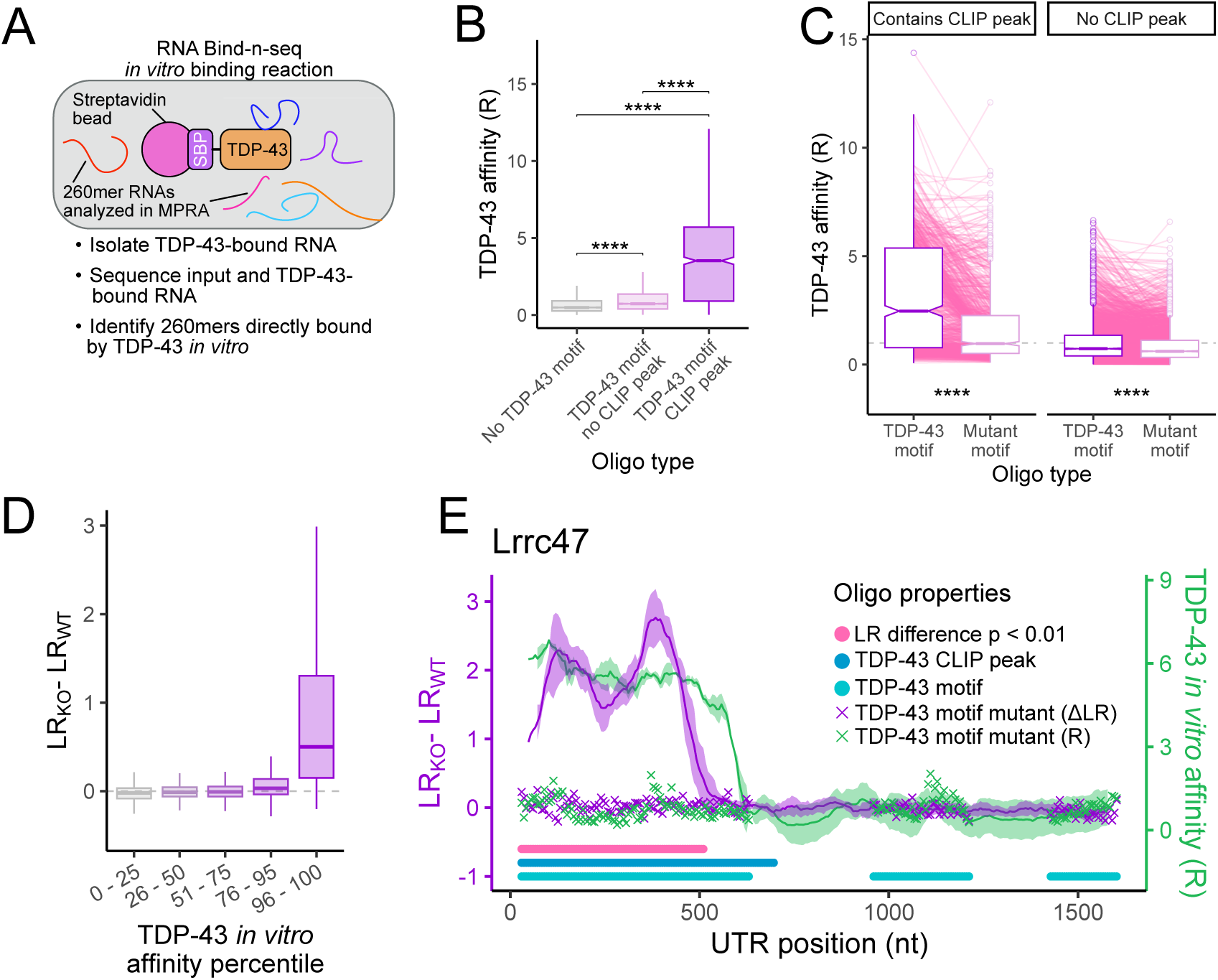
Sequences that regulate RNA localization via TDP-43 are directly bound by TDP-43 *in vitro*. (A) Schematic of RNA-Bind-n-seq assay used to determine relative affinities for TDP-43. (B) Differences in TDP-43 affinity for 260mer sequences that do not contain a TDP-43 motif (left), contain a motif but do not lie within a TDP-43 CLIP-seq peak (middle), or contain a motif and lie within an CLIP-seq peak (right). (C) Differences in TDP-43 affinity for RNA sequences that contain TDP-43 motifs and their companion sequences in which those motifs were mutated. (D) Differences in RNA localization to neurites in wildtype and knockout cells for oligos that had low TDP-43 affinities (left) and high TDP-43 affinities (right). (E) TDP-43 affinities (green) and differences in neurite enrichment between wildtype and knockout cells (purple) for all oligos that tile across the Lrrc47 3′ UTR. Oligos that contain a TDP-43 motif are represented by a light blue circle, those that contain a TDP-43 CLIP-seq peak are represented by a dark blue circle, and values for oligos with mutated TDP-43 motifs are represented by a purple and green x marks. P values were calculated using Wilcoxon ranksum tests. NS (not significant) represents p >0.05, * p < 0.05, ** p < 0.01, *** p < 0.001, and **** represents p < 0.0001.

Mirroring the MPRA data, we found that while a motif alone was sufficient to give a slight increase in affinity, the increase in affinity was much larger if that motif lied within a TDP-43 CLIP-seq peak (**Figure 5B**). Mutation of the TDP-43 motif within an oligo drastically reduced its affinity for TDP-43, again much more so if that motif laid within a TDP-43 CLIP-seq peak (**Figure 5C**). These results reinforce the idea that the information that discriminates between occupied and non-occupied TDP-43 motifs is contained within the local sequence context. They also demonstrate that the motif discrimination happening in cells can be recapitulated *in vitro* [48].

To more directly compare the MPRA and RBNS results, we compared LR (function) and R (binding) values for each oligo. We found that the oligos with the highest R values also had the highest differences in LR values between wildtype and knockout cells (**Figure 5D**), indicating that TDP-43-dependent RNA localization activity in cells can be predicted from *in vitro* affinity. Similarly, oligos with the highest LR differences had the highest R values, indicating that TDP-43 affinity can be predicted from RNA localization activity (**Figure S4C**).

These results can be summarized by visualizing LR and R values for oligos drawn from across a single 3′ UTR (**Figure 5E**). For the Lrrc47 3′ UTR, oligos from the 5′ end of the UTR show both a high affinity for TDP-43 *in vitro* (green line) and sufficiency to cause a reporter RNA to be more neurite-enriched when TDP-43 is lost (purple line). These oligos overlap with a TDP-43 CLIP-seq peak (dark blue line), and mutations of the TDP-43 motifs within these oligos abrogate both the affinity of these oligos for TDP-43 and their ability to regulate RNA localization in a TDP-43-dependent manner. In contrast, although oligos from the the 3′ end of the UTR contain TDP-43 motifs (light blue line), they are not sufficient to regulate RNA localization in a TDP-43-dependent manner and are not bound by TDP-43 in cells or *in vitro*.

### TDP-43 regulates RNA localization by promoting RNA instability

We found that the _TDP-43 represses the_ neurite localization of specific RNAs, and that it does so by binding discrete motifs in their 3′ UTRs. However, the mechanism by which TDP-43 regulates RNA localization was unclear. A recent study reported that changes in the stability of an RNA can have predictable effects on the localization of an RNA in neurons with an increase in stability for an RNA resulting in an increased abundance of that RNA in neurites [49]. Given that TDP-43 is known to promote RNA instability [30,50], we reasoned that TDP-43 loss may result in an increased stability of its RNA targets, leading to their observed increase in neurite abundance.

To test this hypothesis, we used SLAM-seq to measure the stability of the same group of thousands of reporter RNAs used in the RNA localization MPRA (**Figure 4A**) in wildtype and TDP-43 knockout cells. SLAM-seq uses pulse-chase metabolic labeling of cellular RNA with 4-thiouridine (4SU) [51]. At given chase timepoints following the 4SU pulse, RNA is collected and 4SU within cellular RNA is quantified by high-throughput sequencing as T>C conversions to determine relative stabilities for all RNAs in the sample. As in the MPRA, we performed targeted RNA sequencing of our pool of reporter-embedded oligos. We then compared their 4SU content in wildtype and knockout cells at 0 hour and 12 hour uridine chase timepoints (**Figure 6A**).

**Figure 6.**
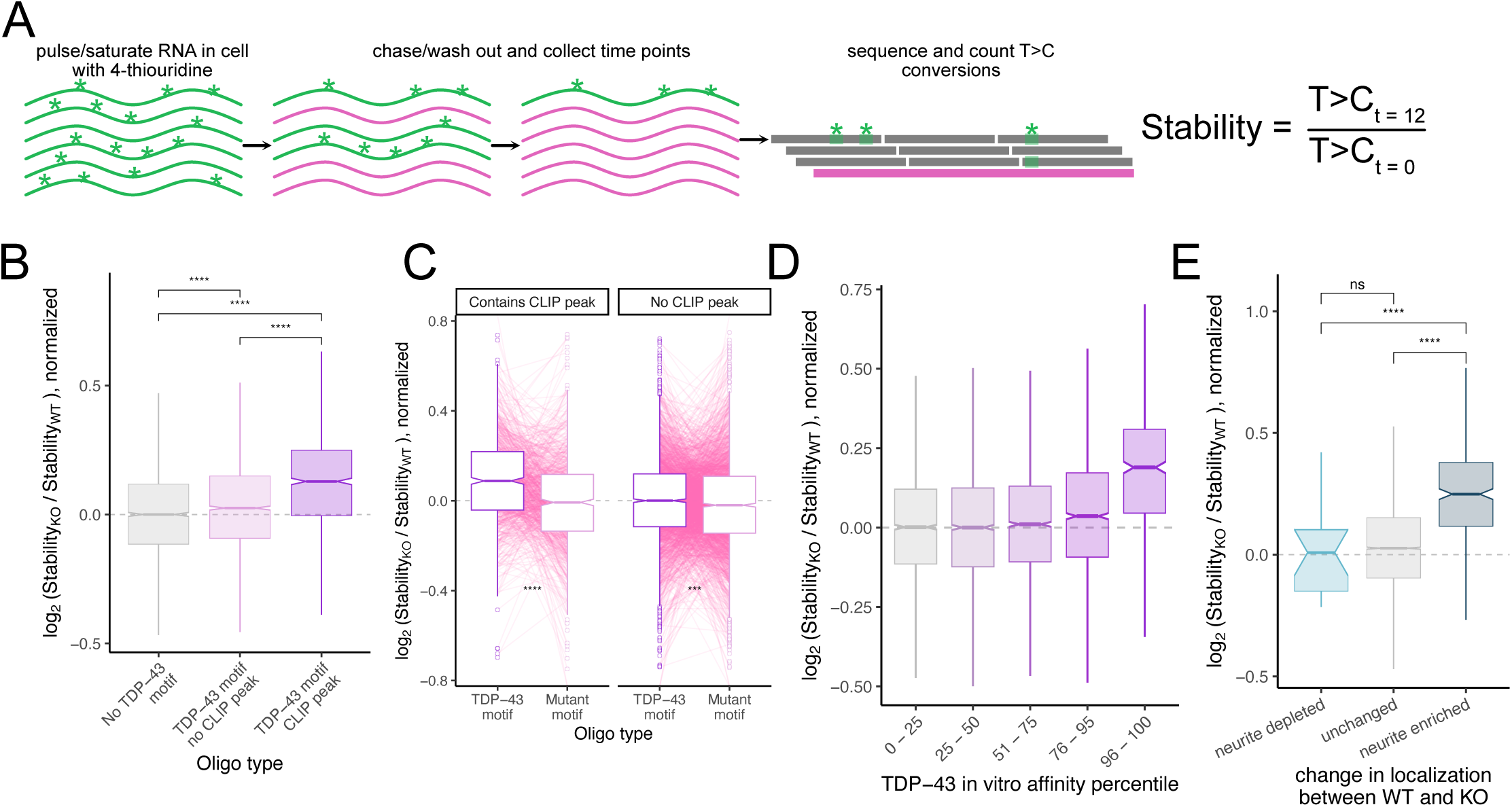
TDP-43 regulates RNA localization through regulating RNA stability. (A) Schematic of SLAM-seq experiment for the study of RNA stability. RNA samples were collected at 0 hr and 12 hr following the chase with unlabeled uridine. For each oligo, its relative stability in a given sample was calculated by comparing rates of T>C conversions in the 12 hour and 0 hour timepoints. (B) Differences in stability between wildtype and knockout cells for reporter RNAs containing oligos that do not have a TDP-43 motif (left), contain a motif but do not lie within a CLIP-seq peak (middle), or contain a motif and lie within a CLIP-seq peak (right). (C) Changes in stability between wildtype and knockout cells for reporter RNAs with oligos that contain TDP-43 motifs and their companion oligos in which those motifs were mutated. (D) Changes in stability between wildtype and knockout cells for reporter RNAs with oligos that showed low affinity for TDP-43 in vitro (left) and those that showed high affinity for TDP-43 (right). (E) Changes in stability between wildtype and knockout cells for reporter RNAs that showed the indicated change in neurite localization between wildtype and knockout cells. P values were calculated using Wilcoxon ranksum tests. NS (not significant) represents p >0.05, * p < 0.05, ** p < 0.01, *** p < 0.001, and **** represents p < 0.0001.

To test our ability to perform SLAM-seq on a pool of thousands of reporters, we performed a small pilot experiment. We found that, as expected, our MPRA reporters only contained T>C conversions when both 4SU was added to cells and RNA was treated with the alkylating agent iodoacetamide (IAA) (**Figure S5A**). We then moved to calculating the stability of our reporters within and without TDP-43. To do this, we treated cells with 4SU for 24 hours and chased with unlabeled uridine for 12 hours. We then calculated relative stabilities for all reporters by comparing T>C conversion rates at the beginning of the chase period (t = 0) to those after 12 hours of chase (t = 12). To quantify how a reporter’s stability depended on TDP-43, we then asked how these t = 12 / t = 0 stability ratios differed between wildtype and knockout samples (**Table S5**).

Again mirroring the MPRA and RBNS results, reporter-embedded oligos that contained TDP-43 motifs showed significantly increased stability upon TDP-43 loss, consistent with TDP-43 promoting RNA decay. However, the effect size of this increased stability was much larger if that oligo lied within a TDP-43 CLIP-seq peak (**Figure 6B**). As with the MPRA and RBNS, mutation of the TDP-43 motifs abrogated this effect (**Figure 6C**). Similarly, oligos that had the highest affinity for TDP-43 in the RBNS assay showed the largest increases in stability upon TDP-43 loss (**Figure 6D**). Finally, oligos that mediated increased neurite enrichment upon TDP-43 loss in the MRPA also showed the largest increases in stability upon TDP-43 loss (**Figure 6E**). These results connect TDP-43 affinity for unstructured motifs, TDP-43’s ability to modulate RNA stability, and TDP-43-mediated changes in RNA localization. Together, they support a mechanism where RNAs with unstructured TDP-43 motifs are bound by TDP-43 and destabilized. When TDP-43 is knocked out, that repression is released, allowing more of those RNAs to survive long enough to complete the journey to neurites.

### Evidence of loss of TDP-43-mediated regulation of RNA stability in patient samples

In ALS samples, TDP-43 is often aggregated, leading to a loss of function [52]. We wondered whether previously reported changes in RNA stability in ALS patient samples could be linked to signatures of the loss of TDP-43 activity. We used previously reported data in which RNA stability had been measured transcriptome-wide in iPSCs derived from sporadic ALS (sALS) and C9orf72-mediated ALS (C9ALS) using BruChase-seq [31]. We compared the stability of RNAs in these samples to those from unaffected patient controls. For both sALS and C9ALS, we found a significant positive correlation between the number of TDP-43 motifs in the 3′ UTRs of RNAs and their increase in stability in ALS samples compared to control. In both cases, RNAs with the highest number of TDP-43 motifs in their 3′ UTRs showed the highest increase in stability in disease samples (**Figure S5B,C**).

TDP-43 can also be induced to aggregate by overexpressing it without its self-regulatory endogenous 3′ UTR [53,54]. When we looked in previously published stability data from iPSC cells in which TDP-43 had been overexpressed [31], we again found that RNAs with the most TDP-43 motifs in their 3′ UTRs showed the greatest increase in stability upon TDP-43 overexpression (**Figure S5D**). These results suggest that loss of TDP-43 function in ALS results in the increased stability of its RNA targets. Given our results linking TDP-43-mediated stability and RNA localization, they further suggest that TDP-43 target RNAs may be mislocalized in ALS samples.

### TDP-43 inhibits RNA localization to axons in primary mouse neurons

As we had established that TDP-43 binding and changes in RNA stability lead to changes in RNA localization in our mouse neuronal cell line, we then asked if these patterns are observed in other systems. One published study reported changes in soma and axon RNA content following TDP-43 knockdown in primary mouse dorsal root ganglia [34]. When we compared this data to our own, we found that the RNAs that showed increased neurite enrichment in TDP-43 knockout CAD neuronal cells also showed axon enrichment following TDP-43 knockdown in the primary DRG data (**Figure 7A**). Further, the RNAs that became more axon-enriched after TDP-43 knockdown in the DRG data contained significantly more TDP-43 motifs in their 3′ UTRs than expected (**Figure 7B**). These results suggest that CAD cells are generally faithful reporters of RNA localization mechanisms that operate in primary cells, and that, as in CAD cells, TDP-43 binds 3′ UTRs to repress RNA accumulation in the axons of primary neurons.

**Figure 7.**
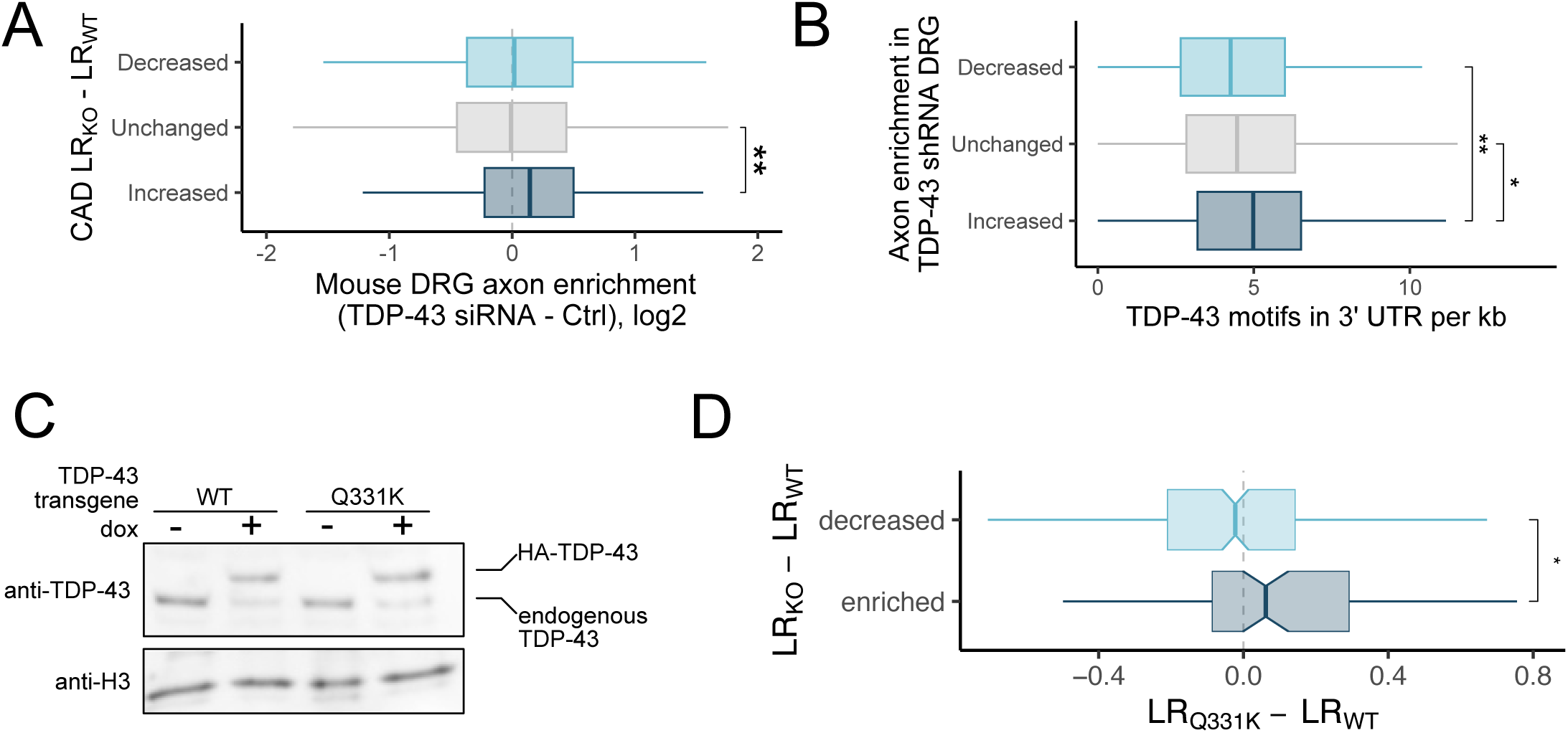
Evidence of TDP-43-mediated RNA localization in other neuronal systems and of its misregulation in ALS samples. (A) Changes in axon enrichment in mouse DRG samples following TDP-43 knockdown for RNAs with the indicated changes in localization following TDP-43 knockout in CAD cells. (B) TDP-43 motif density in the 3′ UTRs of RNAs with the indicated changes in localization following TDP-43 knockdown in mouse DRG. (C) Immunoblot of CAD cells with single copy, site-directed integration of doxycycline-inducible wildtype or Q331K TDP-43 transgenic alleles. (D) Changes in neurite RNA localization in cells expressing Q331K TDP-43 compared to wildtype TDP-43 for RNAs with the indicated changes in localization following TDP-43 knockout in CAD cells. P values were calculated using Wilcoxon ranksum tests. NS (not significant) represents p >0.05, * p < 0.05, ** p < 0.01, *** p < 0.001, and **** represents p < 0.0001.

### TDP-43 inhibits RNA localization to neurites in human motor neurons

To further probe RNA localization mechanisms in more physiologically relevant systems, we differentiated human motor neurons from iPSCs and quantified their RNA localization patterns by fractionating them into soma and neurite fractions as before [55] (**Figure 1A**). To quantify the effect of TDP-43 on RNA localization in this system, we performed this experiment after treating cells with an shRNA targeting TDP-43 or a scrambled control shRNA (**Figure S6A,B**). We found that RNAs that became more neurite-enriched following TDP-43 knockdown had significantly more TDP-43 motifs in their 3′ UTRs than those than expected, again consistent with TDP-43 generally acting to keep RNAs out of neurites (**Figure S6C**).

### ALS-associated mutations in TDP-43 inhibit its ability to regulate RNA localization

Given that mutations in TDP-43 are strongly associated with ALS, we decided to investigate what effects they may have on RNA localization. To do this, we took advantage of TDP-43’s autoregulatory activity to create CAD lines in which the majority of the expressed TDP-43 was either wildtype or contained an ALS-associated mutation (Q331K). Using cre-lox recombination, we site specifically integrated a single-copy of wildtype or mutant, HA-tagged TDP-43 with a dox-inducible promoter and synthetic 3′ UTR into wildtype CAD cells. Because TDP-43 autoregulates its own expression by binding its 3′ UTR to induce degradation of its RNA [50], dox-induced expression of the transgene resulted in downregulation of the endogenous alleles (**Figure 7C**). The transgene was immune from this regulation as it contained a synthetic 3′ UTR. However, because the transgene is integrated as a single copy, the overall level of TDP-43 remained approximately equal in induced and uninduced cells (**Figure 7C**). Thus, we were able to engineer lines in which we could avoid the overexpression of TDP-43 that can lead to adverse effects [53,54] but in which the majority of TDP-43 in the cell was either wildtype or Q331K.

We then fractionated these cells to assay changes in RNA localization between cells expressing wildtype and Q331K TDP-43. In general, the RNAs that became more neurite-enriched in cells expressing Q331K TDP-43 were the same RNAs that became more neurite-enriched in TDP-43 knockout cells (**Figure 7D**). This suggests that Q331K TDP-43 is a loss-of-function mutation in the context of RNA localization and that RNAs containing sites of TDP-43 interaction in their 3′ UTRs may be mislocalized in ALS patient samples.

## DISCUSSION

Most of the work done on TDP-43’s impact on RNA metabolism has focused on its ability to regulate alternative pre-mRNA splicing and RNA decay [31,56,57]. This work has identified connections between TDP-43 regulation of specific RNAs and ALS phenotypes [29,56]. However, as with many RBPs, TDP-43 regulates hundreds of RNAs at multiple points in RNA metabolism, raising the possibility that observed phenotypes may be due to many points of misregulation.

Those studies that have focused on the regulation of RNA trafficking by TDP-43 have generally concluded that TDP-43 promotes RNA transport to neuronal projections. The evidence supporting these conclusions has included the fact that TDP-43 is transported along axons [35] and that knockdown of TDP-43 resulted in decreased transport of RNP granules [33]. However, from these studies it was unclear which RNAs, if any, were mislocalized following TDP-43 perturbation. Other studies have found that axonal transcriptomes contain different RNA populations upon TDP-43 perturbation [34,36], but it was unclear which of these changes were directly due to the loss of TDP-43 binding on the RNA and which were due to secondary effects of TDP-43 loss.

Here, using a variety of high-throughput and targeted assays, we have found that TDP-43 directly acts to inhibit RNA accumulation in neurites and likely does so through its ability to negatively regulate RNA stability. This is consistent with previous reports of other RBPs that negatively regulate RNA stability also acting to prevent neurite accumulation of their targets [17,49,58]. In the course of these experiments, we tested TDP-43’s ability to regulate the localization of thousands of discretely defined sequences drawn from endogenous 3′ UTRs using a massively parallel reporter assay. We then tested TDP-43’s ability to bind and regulate the stability of those exact same sequences using RNA Bind-n-seq and SLAM-seq, respectively. Because we performed all three of these assays on the same pool of RNA sequences, we were able to make inferences about the interconnectedness of the three processes. We envision this approach as a powerful one for future studies of RBP activity.

Clear links have been made between the misregulated splicing of specific transcripts that arise due to TDP-43 loss of function and ALS phenotypes [29,56]. However, the contribution that RNA mislocalization makes to ALS phenotypes, if any, is still unknown. Although mutations in a number of RBPs that regulate RNA localization are associated with a variety of neurologic diseases [23], it has been difficult in mammals to assign phenotypic consequences to the mislocalization of specific RNAs. By identifying cis-elements within RNAs that regulate their localization through the activity of specific RBPs, we hope that the work presented here will inform future experiments that will perturb the trafficking of specific RNAs, allowing the formation of mechanistic connections between RNA transport and neurological phenotypes.

## ACKNOWLEDGEMENTS

We thank members of the Taliaferro lab and Wilfried Rossoll for helpful discussions and guidance. This work was funded by NIH grants R35GM133385 (JMT), R01NS122911 (JMT and HAR), R35GM142864 (DD), F31GM151819 (CM), F31GM155957 (KV), and T32GM136444 (CM).

## METHODS

### Cell culture

CAD cells were grown in DMEM/F-12 (Gibco, #11320-033) supplemented with 10% Equafetal (Atlas Biologicals, #EF-0500-A) and 1% Penicillin–Streptomycin solution (Gibco, #15140122). The cells were grown in a humidified incubator at 37°C and 5% CO2.

### Creation of TDP-43 inducible knockout cell lines

Since TDP-43 is an essential protein in many cell lines, we devised a strategy to generate doxycycline sensitive conditional knockout TDP-43 CAD and N2A cell lines. First, we designed a HA-tagged TDP-43 transgene with multiple silent mutations in two sgRNA binding sites and PAM sites, such that the transgene is immune to gRNA cutting. In addition, the gRNA-resistant TDP-43 ORF was followed by T2A and Neomycin resistance gene, ending with 3′-UTR of TDP-43. This transgene was cloned into a modified version of pRD-RIPE [40] that was changed to be dox-off instead of dox-on. Cells were co-transfected with this plasmid mixed with 1% of plasmid expressing Cre recombinase. To transfect one well of a 6-well plate, 1.5 µg of reporter plasmid and 15 ng of Cre-plasmid was mixed with 3 µl Lipofectamine LTX reagent (Invitrogen, #15338100), 1.5 µl PLUS reagent and 100 µl Opti-MEM following the manufacturer’s protocol. These cells were incubated with the transfection mixtures for 24hr. Media was then changed and the cells were allowed to rest for another 24 hours. They were then selected with 2.5µg/mL puromycin to generate stable lines.

CAD and N2A lines with the gRNA resistant HA-tagged TDP-43-T2A-Neo transgene stably integrated were then treated with sgRNAs to target endogenous TDP-43 alleles. Two sgRNAs that target exon2 and exon6, respectively (ACCAUCAGAAGACGAUGGGA and CUCCACCCAUAUUACCACCC) were obtained from Synthego and dissolved in 1× TE buffer at a final concentration of 100 µM. Purified Cas9 protein was also obtained from Synthego. RNP complexes were assembled using a 3:1 molar ratio of sgRNA:Cas9 at room temperature for 15 min. The RNP complexes were then co-electroporated with a plasmid encoding GFP into CAD cells using the Neon transfection system (ThermoFisher) using the following settings: 1400 V, 1 pulse for 30 ms. The cells were allowed to recover for 72 hours and then GFP-positive single cells were isolated using a MoFlo XDP100 cell sorter. The single cells were allowed to grow undisturbed for 2 weeks in the presence of G418 (800 µg/ml) and then expanded for screening.

### Screening of CRISPR clones

Genomic DNA from single cell clones was isolated using the PureLink Genomic DNA Mini kit (Invitrogen, #K182002). The clones were initially screened for the deletion of a portion of the TDP-43 coding sequence by PCR using primers flanking the gRNA target sites in exon 2 and exon 6.

Integration of the TDP-43 transgene and confirmation of the KO was assayed by western blot by using antibodies against TDP-43 (Proteintech, #10782-2-AP, 1:10,000 dilution), HA tag (transgene, GenScript, #A01244, 1:10,000 dilution), and histone H3 (loading control, Abcam, #10799, 1:10,000 dilution).

KO cells are maintained in DMEM/F-12 with 10% Equalfetal (Atlas Biologicals) and 1% Penicillin–Streptomycin solution supplemented with 2.5µg/mL puromycin.

Lastly, once clones with no endogenous expression of TDP-43 were selected, the TDP-43-T2A-Neo transgene cassette was switched using RMCE with a similar plasmid. This plasmid contained a blasticidin resistance gene instead of puromycin resistance and contained the same TDP-43 transgene and its endogenous 3’ UTR but lacked the T2A-Neo resistance gene. The cells were selected in blasticidin (5 µg/ml) and expanded in blasticidin.

### Subcellular fractionation into soma and neurite samples

All subcellular fractionations were performed as in [39]. Briefly, cells were plated at 90% confluency in complete media on transwell membranes with 1 µm pores (Corning, #353102). Cells were allowed to attach for 1 hour, then media was replaced with serum-free media or serum-free media with 1 µg/mL doxycycline to turn off TDP-43 expression. 48 hours later, cells were fractionated by scraping the top of the membranes with a cell scraper to remove soma while neurites remained stuck to the underside of the membrane. RNA was then isolated from each fraction using a Quick RNA Micro Kit (Zymo, #R1051).

### Sequencing of endogenous RNA samples

For the sequencing of endogenous RNA samples, 100 ng of total RNA was prepared into RNAseq libraries using the KAPA mRNA Hyperprep library kit (Roche, #08098123702). Libraries were amplified using 16 PCR cycles.

### Analysis of soma and neurite endogenous RNA samples

Read sequences were trimmed of adapters using cutadapt [59]. Transcript abundances were calculated using salmon [60], and gene-level abundances were quantified using tximport [61]. Reads were quantified against the mm10 genome and Gencode version 17. Differentially abundant RNAs between soma and neurite fractions were identified using DESeq2 [62]. RNAs with different neurite enrichments between genotypes were identified using xtail [63].

### Analysis of neurite lengths

To measure neurite length of CAD TDP43 DOX-OFF cells with +/- dox treatment, ∼200,000 cells were seeded into each well of a CellTreat 6-well tissue culture plate with 22 mm Poly-D-lysine coated coverslips (Neuvitro, CAT# H-22-1.5-PDL) placed within the wells. Cells were incubated at 37C with 5% CO2 for 1 hour in complete F12:DMEM growth media, afyter 1 hour, growth media was then replaced with serum free F12:DMEM media for 48 hours to induce neurite differentiation and +/- dox treatment to turn off TDP-43 expression. 48 hours post differentiation, cells were washed with PBS -/- then fixed with 3.7% formaldehyde in PBS -/- for 15 minutes. After fixation cells were washed with PBS -/- then permeabilized with 0.1% Triton-X100 for 5 minutes, followed by washing the cells with PBS -/-. After permeabilization, cells were blocked with 3% BSA (Fisher Scientific, CAT# 1265925GM) in PBST 0.05% for 30 minutes at room temperature. Cells were then incubated with the Mouse monoclonal anti-Kif5a antibody (1:1000 dilution, Millipore-Sigma, CAT# MAB1614) in blocking buffer overnight at 4°C. Kif5a antibodies were used since this motor protein provides clear visualization of the complete neurite length. Cells were then washed 3 times with PBST 0.05% for 5 minute increments. After washing, cells were incubated with the anti-mouse AF488-conjugated Fab fragment (1:1000 dilution, Cell Signaling technologies, CAT# 4408S) secondary antibody in blocking buffer for 1 hour at room temperature. Cells were then washed again with PBST 0.05% 3 times for 5 minute increments and maintained in PBS -/- until mounting. To mount the coverslips to the coverglass (Globe Scientific, CAT# 1380-20), immunostained coverslips were mounted with 20 uL Fluoromount G (Southern biotech, CAT# 0100-01) for 20 minutes at room temperature. Coverslips were secured to the coverglass using nail polish on 4 regions of the coverslip for 30 minutes at room temperature. Immunostained cells were imaged on the EVOS M7000 widefield benchtop microscope with the 10x objective using the GFP (470/525 nm) filter set. Three fields of view were imaged per +/- dox treatment. Fields of view were saved as TIFF files to retain pixel sizes and proper image formatting for accurate micron distances.

To analyze +/- dox treated anti-Kif5a labeled neurite lengths, the open-source ICY bioanalysis software was utilized [64]. EVOS TIFF files were loaded onto ICY and neurite lengths were analyzed using the polyline 2D ROI detection tool to manually measure the length of neurites (base of the neurite/soma region to the end of the neurite synapse) within the image field. The polyline 2D ROI detection method allows the user to accurately draw a 2D line with multiple nodes to allow accurate detection of neurite lengths. Long neurites (>20 um) were quantified with the polyline 2D ROI tool with. Cells with multiple neurites were also analyzed, but soma projections/dendrite-like projections were not quantified (most cells have only 1 neurite). To derive the neurite lengths from the ROIs, the ROI statistics tool was used with the perimeter function to derive the micron distance per neurite analyzed. Neurite lengths were statistically measured using the Wilcoxon non-parametric test.

### Visualization of RNA localization using single molecule RNA FISH

CAD TDP-43 KO cells were plated on PDL coated glass coverslips (neuVitro, H-18-1.5-PDL) that fit within 12 well plates at 4.0 x 104 cells per well in full growth media or full growth media with 2.0µg/mL doxycycline for 3 hours to allow cells to attach to the coverslip. Media was removed and replaced with serum free media or serum free media with 2.0µg/mL doxycycline to induce neurite differentiation for 48 hours. The media was aspirated, and cells were washed once with 1× PBS. Cells were fixed in 3.7% formaldehyde (Fisher Scientific, #BP531-500) in PBS for 10 min at room temperature and then washed twice with 1× PBS. Cells were permeabilized with cold 70% ethanol (VWR, #EM-EX0276-3S) for 2 hours at 4°C. The cells were washed with freshly prepared wash buffer (2X SSC (Fisher Scientific, #15557044) and 10% formamide (Sigma-Aldrich, #F7503-100ML) in water) at room temperature for 15 minutes. In the meantime, the smiFISH probes for each gene were hybridized to the fluorescent Y Flap using the protocol from Tsanov et al. (2016)[42]. Diras1 and Ksr2 probe sets each included 48 probes.

After hybridization, the probe/flap hybridization product was spun in a benchtop centrifuge for 60 seconds. Per coverslip, 2 µl (0.833 µM) of smFISH probe was added to 100 µl of smFISH hybridization buffer (Biosearch Technologies, SMF-HB1-10) for each condition and gene. A hybridization chamber was prepared using an empty tip container, wrapped in tinfoil with parafilm and wet paper towels inside the box to retain moisture. 100 µl of the probe-containing hybridization solution was added to the parafilm. The coverslip was then placed on top of this droplet of hybridization buffer with the cell side down. The hybridization chamber with the coverslips was incubated at 37°C overnight (15-18 hours). The coverslips were transferred to a fresh 12-well plate with the cell side up and incubated twice with freshly prepared wash buffer for 45 minutes at 37°C. The second wash included a membrane dye (CellBrite® Fix 488, Biotium, #30090-T). Then slides were incubated with wash buffer including DAPI (100 ng/mL) at 37°C for 30 minutes. Slides were washed twice with PBS for 5 minutes at room temperature. Coverslips were then mounted onto slides with Fluoromount G (SouthernBiotech, #0100-01) and sealed with nail polish.

Slides were imaged at 63x with consistent laser intensity and exposure times across samples. DAPI was imaged with an exposure of 100ms. Membrane dye was used to find cells and was imaged in the FITC channel with an exposure of 100ms. FISH probes were visualized in the TRITC channel with an exposure of 1500ms. Z stacks were collected of approximately 24 images, 0.4µm apart.

### Analysis of smFISH data

FISH-quant was used to quantify neuronal projection enrichment of smFISH spots as previously described [9]. Briefly, outlines were drawn in the FITC channel visualizing the cell membrane. Two outlines were drawn per cell: soma and neuronal projection. Prior to quantification, identified smFISH spots were thresholded for intensity, sphericity, amplitude, and position. Transcript enrichment was quantified by the total number of spots in the projection of cells over a total number of spots in the soma.

### Assaying RNA localization of single defined reporter transcripts via RT-qPCR

Three wells of the 6-well plate served as one replicate for the RT-qPCR experiments. Cells were transfected, plated, and fractionated as previously described. 100ng of RNA from each soma and neurite sample were reverse transcribed using LunaScript RT SuperMix (New England Biolabs, #E3010) in a 10µL reaction volume following manufacturer’s instructions. The resulting cDNA was then diluted to 20µL total volume using nuclease-free water. 2µL of diluted cDNA is used as the template for qPCR to estimate the abundances of Firefly and Renilla reporter transcripts in the soma and neurite fractions. The qPCR reaction was performed using PrimeTime® Gene Expression Master Mix (Integrated DNA Technologies, #1055772) with differently labeled probe sets for each pair of transcripts, allowing the quantification of two transcripts in the same reaction. Firefly probe sets (Integrated DNA Technologies) are HEX labeled while Renilla probe sets (Integrated DNA Technologies) are FAM labeled.

The observed Ct values of the transcripts were within the recommended dynamic range of the assay. Reactions were carried out using a CFX-Opus 384 thermocycler with the following conditions: UNG activation at 50°C for 2 min, followed by polymerase activation at 95°C for 30 s and 40 cycles of 95°C for 5 s, and 60°C for 30 s. Finally, a melting curve was performed by incubating samples at 65°C for 15 s followed by a temperature gradient increase at 0.5°C/s to 95°C. Each sample was measured with three technical replicates. To ensure no contamination, no reverse transcriptase and no template controls were performed. Fold enrichment was calculated using the ΔΔCt method. MIQE guidelines were followed for all qPCR experiments.

### Oligonucleotide design for MPRA

The code for designing the MPRA oligos is available at https://github.com/TaliaferroLab/OligoPools/blob/master/makeoligopools/OligoPools_shortstep_260nt.py. This script designed 260 nt oligonucleotides with a step size in between neighboring oligonucleotides of 6 nt. For a given gene, oligonucleotides were designed against the 3′ UTRs of all protein-coding transcripts with well-defined 3′ ends (as defined as not having the ‘cds_end_NF’ or ‘mRNA_end_NF’ tags). The polyA site of the UTR was also required to be positionally conserved in humans. This was assessed by getting the syntenic region surrounding the mouse polyA site in the human genome using UCSC liftOver. A polyA site was defined as conserved if there was a polyA site within 200 nt of the corresponding region of the human genome. Additionally, UTRs longer than 10 kb were excluded. Any oligo with a TDP-43 motif (defined as GUGUG, UGUGU, or GUAUG) also had a companion oligo in which those motifs were mutated (to CACAC, ACACA, and CAUAC, respectively)

UTRs of multiple filter-passing transcripts for a single gene were merged together to create a meta-UTR. Oligonucleotides were then designed against this meta-UTR with the addition of 260 nt upstream and downstream of the beginning and end of the UTR in order to give full coverage of the ends of the UTR with multiple oligonucleotides. 20 nt PCR handles were then added to the ends of every oligonucleotide. The pool of oligonucleotides was synthesized by Twist Biosciences.

### Creation of MPRA plasmid pool

The oligonucleotide pool containing the MPRA oligos was obtained from Twist Bioscience and resuspended in 10 mM Tris–EDTA buffer, pH 8.0 to a concentration of 10 ng/µl. The pool was amplified by performing 8 PCR reactions 50 µl each, using Kapa HiFi HotStart DNA Polymerase (Kapa Biosystems, #KK2601) according to the manufacturer’s instructions. A total of 20 ng of the original pool was used as the input DNA template in the 8 PCR reactions with 15 amplification cycles. Following amplification, the PCR reaction was treated with Exonuclease I, at 37°C for 2 h to digest the single-stranded template and primers. The DNA was then purified using 0.8X magnetic beads from Axygen (AxyPrep MAG PCR Clean-Up, #MAG-PCR-CL-50) according to the manufacturer’s protocol.

The pCMV-GFP reporter plasmid was linearized by digesting it with BstXI (NEB, #R0113) at 37°C for 4 h to clone the library into the 3′-UTR of GFP reporter. Digested plasmid DNA was gel extracted using Zymoclean Gel DNA Recovery Kit (Zymo Research, #D4008). The digested plasmid and amplified DNA library were ligated using Gibson Assembly reaction (NEB) using the insert:vector molar ratio of 4:1 at 50°C for 1 hour. The cloned reporter plasmid (∼200 ng DNA) was ethanol precipitated to get rid of excess salts and was then transformed into Escherichia coli using MegaX DH10B T1R Electrocompetent Cells (ThermoFisher, #C640003), using a Biorad GenePulser electroporator. The transformed cells were grown in recovery medium at 37°C for an hour and then plated on pre-warmed Luria broth (LB) agar-Carbenicillin 15-cm plates and incubated at 37°C overnight. The next day, the colonies were harvested from the plates using LB medium and spreader. The bacterial culture was then centrifuged at 4000 rpm for 20 min. The reporter plasmids libraries were purified using ZymoPURE Plasmid Maxiprep kit (Zymo Research, #D4203). Restriction digestion was performed to confirm that the plasmid library contains only a single insert of the right size.

### Assaying TDP-43-dependent RNA localization via MPRA

Cells for MPRA were transfected at 80% confluence in a T-75 flask with 15µg of the MPRA plasmid pool, 48µL Lipofectamine LTX reagent (Invitrogen, #15338100), 24µL PLUS reagent, and 1.2mL of Opti-MEM following the manufacturer’s protocol. Cells were incubated with the transfection mixtures for 2-6 hours, followed by a media change. 24 h later, media was changed again and 1 µg/mL doxycycline was added. 24 h later, cells were plated on transwell membranes and fractionated as in [9].

As in [9], 500 ng total RNA from each soma and neurite fractions was taken to synthesize into cDNA in a 20 µl reaction using SuperScript IV Reverse Transcriptase (ThermoFisher, #18090200) according to the manufacturer’s protocol, with primers specific to Firefly and GFP CDS containing an 8-nt unique molecular identifier (UMI) and a partial Illumina read 1 primer sequence. The incubation time at extension step (55°C) was increased to an hour. Post reverse transcription, 1µL each of RNAse H and RNaseA/T1 mix was added directly into the RT-reaction and incubated at 37°C for 30 min to digest the remaining RNA and RNA:DNA hybrids. The cDNA was purified using Zymo DNA Clean & Concentrator kit (Zymo, #D4013) using 7:1 excess of DNA binding buffer recommended for binding ssDNA.

For library preparation, each purified reporter cDNA reaction was split into five PCR reactions (4 µl cDNA/PCR) and amplified using a reporter specific forward primer with Illumina sequencing adaptors and a reverse primer binding the partial Illumina read 1 sequence with the remaining sequencing adaptors using Kapa HiFi HotStart DNA Polymerase (Kapa Biosystems, #KK2601) using 18× cycles for GFP and 23× for Firefly reporter. The five PCR reactions per sample were pooled together and purified using double SPRI beads protocol. In the first purification round, 0.5× SPRI beads were used to get rid of longer DNA products. The supernatant from this purification was then removed and additional SPRI beads were added to bring the final overall concentration to 0.8×. DNA bound to these beads was then washed and eluted. The library was quantified using Qubit dsDNA HS Assay Kits (ThermoFisher, #Q32854) and size of the library was verified using Tapestation (Agilent D1000 ScreenTape, #5067-5582).

### Analysis of MPRA results

Adapters were removed from reads using cutadapt [59]. Specifically, the sequences GGCGGAAAGATCGCCGTGTAAGTTTGCTTCGATATCCGCATGCTA and CTGATCAGCGGGTTTCACTAGTGCGACCGCAAGAG were trimmed from the 5′ ends of the forward and reverse reads, respectively. The trimmed reads were then aligned to the reference oligonucleotide sequences using bowtie2 and the following parameters: -q –end-to-end –fr –no-discordant –no-unal -p 4 -x Bowtie2Index/ index −1 forreads.fastq −2 revreads.fastq -S sample.sam. Typically, 99% of reads had the expected adapters, and 95% of those aligned to the reference oligonucleotides.

The number of unique UMIs (the first 8 nt of the reverse read) for each reference oligonucleotide was then calculated using https://github.com/TaliaferroLab/OligoPools/blob/master/analyzeresults/UMIsperOligo.py. These UMI counts were then given to DESeq2 [62] to quantify oligonucleotide abundances in each sample and identify oligonucleotides enriched in soma or neurite samples.

### In silico folding of MPRA RNA sequences

RNA sequences were folded using ViennaRNA 2.4.6 [47]. For each 260mer, 80 nt windows were folded at a time, with windows sliding 10 nt across the sequence. For each position, basepair probabilities were then recorded as the summed probability that the base was paired to any other base in the sequence.

### Expression and purification of recombinant TDP-43

Plasmid (pGEX-GST-SBP) containing TDP-43 (residues 1-414, UniProtKB Q13148) was transformed into Rosetta cells (Sigma-Aldrich, #70953) and grown in LB media with 100 mg/mL ampicillin and 25 mg/mL chloramphenicol at 37°C until reaching an optical density of ∼0.8. Cultures were cooled to 4°C and induced with 0.5 mM Isopropyl β-d-1-thiogalactopyranoside (IPTG) for ∼16 hrs at 16°C at which points cells were harvested by centrifugation at 4000 x g for 13 minutes at 4°C. Cell pellets were resuspended in lysis buffer (1% triton X-100, 5 mM DTT, 4 mM MgCl2, 200 mM NaCl, 20 mM HEPES, 1 tablet of Pierce™ protease inhibitor mini tablet, EDTA-free (Thermo Scientific, #A32955) per 2 liters of culture). Lysates were sonicated and subsequently incubated at 25°C for 15 minutes with 250 units of benzonase nuclease (Millipore, #E1014) and 3 units of RQ1 RNAse-Free DNAse per liter of culture. Following nuclease treatment, lysate was centrifuged at 17,500 x g for 30 minutes and supernatant was collected. Supernatant was incubated for 1 hour at 4°C with Glutathione Agarose (Thermo Scientific, #16101) beads to capture GST-tagged TDP-43. Protein-bound beads were washed three times with wash buffer (0.1% triton X-100, 200 mM NaCl, 20 mM HEPES, 3.5 mM EDTA). Protein was eluted with cleavage buffer (10% glycerol, 5 mM DTT, 100 mM NaCl, 20 mM HEPES, 0.01% triton X-100, 1 mg/mL in-house PreScission Protease) for ∼1.5 hours at 25°C. To improve purity, cleaved proteins were passed through a heparin column on a ÄTKA Pure HPLC in a low salt buffer (50 mM NaCl, 50 mM HEPES) and high salt buffer (1 M NaCl, 50 mM HEPES) and relevant fractions were pooled. Triton and glycerol were spiked in for a final concentration of 0.01% and 3%, respectively. Protein was concentrated using an Amicon Ultra 10kDa centrifugal filter unit (#UFC8010) and final concentration and quality was determined by Pierce BCA Assay Kit (Thermo Scientific, #23208) and SDS-PAGE followed by Coomassie staining for visualization.

### Generation of RNA pool for RBNS

Using the plasmid library pool described above we designed primers to amplify inserted fragments and simultaneously add Illumina 5’ and 3’ adapters as well as a T7 promoter. This procedure was done using the following primers:

FWD:TAATACGACTCACTATAGGGAGTTCTACAGTCCGACGATCGCTTCGATATCCGCATGCTA

REV: 5’ CTTGCGGTCGCACTAGTGTGGAATTCTCGGGTGTCAAGG.

PCR cycles were kept to 10-12x and amplification was carried out with Phusion Polymerase (NEB, #M0530) per manufacturer’s specifications. PCR products were resolved on a 2% agarose gel, the product of the correct size excised, and purified using a Qiagen Gel Extraction kit (Cat. #28704). Purified PCR products were used for *in vitro* transcription using the Promega High Capacity T7 In vitro transcription kit (Cat. #P1320) following manufacturers protocols with but with the reaction running for 18hrs at 42°C. RNAs were DNAse (RQ1) treated and purified as previously described [43].

### Quantification of TDP-43 binding to RNA with RNA-bind-n-seq

Recombinant SBP-tagged TDP-43 was diluted to target concentrations in binding buffer (25 mM Tris-HCl, 150 mM KCl, 3 mM MgCl2, 500 mg/mL BSA, 20 units/mL SUPERase IN (Invitrogen)). Dynabeads MyOne Streptavidin T1 (Invitrogen, #65602) beads were washed thoroughly with binding buffer before incubation with recombinant SBP-tagged TDP-43 for 30 minutes at 4°C. Unbound protein was then removed using a magnetic stand (Invitrogen, #12321D) and protein-beads complexes were resuspended with binding buffer. RNA pool with natural TDP-43 targets was added for a final concentration of 500 nM and incubated for 1 hour at 4°C. Following incubation, protein-bound RNAs were washed 3 times with wash buffer (25 mM Tris-HCl, 150 mM KCl, 20 units/mL SUPERase IN (Invitrogen)), washes were performed on a magnetic stand. Samples were eluted twice with elution buffer (4 mM biotin, 25 mM Tris-HCl) for 30 minutes at 37°C and eluates were combined. Eluted RNAs were further purified by phenol chloroform extraction and ethanol precipitation. Isolated RNA was subjected to reverse transcription (SuperScript III Reverse Transcriptase, Invitrogen #18080044) and PCR (Phusion Polymerase, NEB #M0530)) amplified following the same procedures as previously described [43]. RBNS experiments were done in duplicate with two independent protein batches and at 3 different protein concentrations (500, 50, and 5 nM).

### Analysis of RBNS results

RBNS reads were analyzed and assigned to oligos as in the MPRA experiment. Specifically, adapters were trimmed using cutadapt [59], and reads were then assigned to oligo sequences using bowtie2 with the following parameters: -q –end-to-end –fr –no-discordant –no-unal -p 4 -x Bowtie2Index/index −1 forreads.fastq-2 revreads.fastq -S sample.sam. Oligo abundances were then normalized by the total number of mapped reads in each sample, and R values were calculated by dividing normalized abundances in a given sample to those in the input sample.

### Human pluripotent stem cell culture and motor neuron differentiation

Human pluripotent stem cells (hPSC) were cultured on Cultrex ReadyBME (Biotechne, #3434-050-RTU) in mTeSR Plus media (STEMCELL Technologies, #05826) at 37°C in a humidified 5% CO2 atmosphere. hPSCs employed here, have been a derivation from a healthy individual as previously described [65]. For direct differentiation into motor neurons, we used our previously published protocol as described by Hudish et al. with minor modifications [55].

The differentiation was initiated by cluster formation in spinner flasks (ABLE system, ABBWVS03A-6, Reprocell) seeding 0.5e6 cells/ml for 48hrs in mTeSR Plus media in the presence of 10µM Rock Inhibitor Y-27632 (RI, Tocris, #1254). Thereafter, clusters were washed once with KO DMEM (Gibco, #10829018) and transferred to 6 well suspension plates on an orbital shaker (Benchmark Scientific, #BT4001) set at 90 rpm in progenitor medium. Progenitor media consists of base media: KO DMEM, 1:200 Glutamax (Gibco, #35050061), 1:400 N2-A Supplement (STEMCELL Technologies, #07152), 1:200 N21 (Biotechne #AR008), 0.189mM Vitamin C (Sigma, #49752) and 1X Penicillin/streptomycin (Gibco, #15140), that is further supplemented with 1µM Compound C (STEMCELL Technologies, #72102), 10µM SB431542 (STEMCELL Technologies, #72234), 4µM CHIR99021 (STEMCELL Technologies, #72054), and 5µM RI. Clusters were culture for 6 days in this progenitor media with daily partial media changes followed by culture in induction medium for 2 days. Induction media consist of base media supplemented with 1µM Compound C, 10µM SB431542, 200nM SAG (Cayman, #11914), 1.5µM TTNPB (Tocris, #0761) and 5µM RI. Clusters were supplemented with induction media with 10µM RI for day 8-9. At day 9, clusters were dissociated into single cells using 1X TrypLE (Thermo Fisher #12604021) for 12 min at 37°C followed by quenching with base media. Single cell suspensions can be frozen at this time in Cryostor 10 (Biolife Solutions, #210502) at 10e6 cells/ml. Fresh or frozen day 9 progenitors were seeded on transwell cell culture inserts (Corning, #353102) with top and bottom coated with 0.01% poly-l-lysine (Sigma Aldrich, #P4707) and 1xMatrigel (Corning, #354277), respectively, and placed in deep well 6-well cell culture plates (Corning, #353502) in maturation medium comprising of base media containing 10mM glucose, 1x non-essential amino acids (NEAA) (Gibco, #11140), 10µM RI, 200nM SAG, 1.5µM TTNPB, 0.02µg/mL BDNF (STEMCELL Technologies, #78005), and 0.02µg/mL GDNF (STEMCELL Technologies, #78058). Cells are kept in this media for 4 days with daily partial media changes. Thereafter culture media is changed to base media containing 2.5µM gamma-secretase inhibitor (ASIS #0149), 0.02µg/mL BDNF and 0.0µg/mL GDNF, and 2µM RI for 3 days with daily partial media changes. Finally motor neurons (MNs) were cultured in base media containing 0.02µg/mL BDNF and 0.02µg/mL GDNF until day 19.

### Lentiviral transduction, TDP-43 Knockdown and RNA fractionation of human motor neuron samples

Day 11 cells were infected with lentiviral vectors containing a scramble shRNA control sequence (CCTAAGGTTAAGTCGCCCTCG) or an shRNA against TDP-43 (AGATCTTAAGACTGGTCATTC) at 2×105 IU/ml (Infective units/ml) with polybrene (4 µg/mL; VectorBuilder). After 24hrs fresh media was supplemented. Cultures were treated with 0.5µg/ml Puromycin (Thermo, #A1113803) for 4 days with daily media changes to select for infected cells. MNs were harvested at day 19 in antibiotic free maintenance media for the last 24 hours. Mechanical fractionation of cultured neuronal cells into cell body and neurite fractions were performed as described in Arora et al. [39]. RNA was isolated using the Quick RNA Microprep kit (Zymo, #R1051). Six membranes were combined for each single preparation which yielded typically 500ng to 1µg of total RNA. Analysis of knockdown efficiency was checked by western blotting. Total protein was extracted from the cell body suspension using 1x RIPA lysis buffer (Millipore, # 20-188) supplemented with EDTA-free protease inhibitor (Roche, #4693159001) and phosphatase inhibitor (Roche, #04906837001) cocktails. Lysates were resolved on SDS–PAGE, transferred to a nitrocellulose membrane (BioRad, #1620112). After blocking for 60 min, blots were probed with beta-actin (CST, # 4970) and TDP-43 (Proteintech, #10782-2-AP) primary antibodies overnight at 4°C. After washing, membranes were then incubated for 1hr at room temperature with secondary antibody conjugated to horseradish peroxidase (HRP) (Invitrogen, #31460). After incubation with Clarity Western ECL Substrate (BioRad, #1705061), images were captured on a chemiluminescence imaging system (Syngene GeneGnome XRQ).

### Analysis of RNA stability with SLAM-seq

SLAM-seq was performed as described in [51], with a modified RNA isolation procedure. The MPRA plasmid pool was transfected into cells as described above 48 hours before labeling. 24 hours later, cells were seeded at 100% confluency into 6-well plates, allowed to adhere, then subject to differentiation by serum starvation. Cells were labeled with 125µM 4-thiouridine (Sigma-Aldrich, #T4509-25MG) in serum-free media (DMEM/F-12, 1% Penicillin-Streptomycin), replacing media every 12 hours for 24 hours of labeling time, then exchanging media for typical serum-free media. Samples were then collected at time points after exchanging media (0 and 12 hours). Total lysate was collected in RNA lysis buffer and frozen at −80°C for less than 24 hours then isolated using Zymo Quick RNA Microprep kit (Zymo, #1051). All buffers other than the RNA lysis buffer were supplemented with DTT to a final concentration of 0.1mM. Absolute ethanol was supplemented with 0.2mM DTT. Labeled RNA was eluted in water supplemented with 1mM DTT. The optional on-column DNase treatment was used. Eluted RNA was subjected to an additional DNase treatment (New England Biolabs, #M0303) and cleaned up with DTT-supplemented Zymo Quick RNA Microprep kit (Zymo, #1051), starting with the addition of 300µL RNA lysis buffer and skipping the on-column DNase treatment. From each sample, 500ng of alkylated RNA was then subject to targeted RNA library preparation and sequencing as previously described [9].

### Analysis of SLAM-seq results

As with the MPRA, SLAM-seq reads were first trimmed of adapters using cutadapt [59]. Reads were then aligned to oligonucleotide sequences using bowtie2 and the following parameters: -q –end-to-end –fr –no-discordant –no-unal -p 4 -x Bowtie2Index/index −1 forreads.fastq −2 revreads.fastq -S sample.sam. Nucleotide conversions present in each were calculated and assigned to oligonucleotides using a modified version of PIGPEN software [66] available here: https://github.com/TaliaferroLab/OINC-seq/blob/master/getmismatches_MPRA.py.

### Creation of CAD cells expressing wildtype or Q331K TDP-43

CAD cells containing a single loxP-flanked cassette [40] were co-transfected with a plasmid containing HA-TDP-43 transgenes mixed with 1% of plasmid expressing Cre recombinase. To transfect one well of a 6-well plate, 1.5 µg of reporter plasmid and 15 ng of Cre-plasmid was mixed with 3 µl Lipofectamine LTX reagent (Invitrogen, #15338100), 1.5 µl PLUS reagent and 100 µl Opti-MEM following the manufacturer’s protocol. Cells were incubated with the transfection mixtures for 24 hours, followed by the media change. The cells were incubated for an additional 24 hours allowing for recovery and expression of the antibiotic resistance before addition of puromycin (2.5 µg/ml). The cells were selected in the puromycin until the cells in the control wells died. The cells with stably integrated reporter plasmids were expanded in the growth medium with puromycin. Integration was tested by inducing transgene expression with 1 µg/ml doxycycline for 7 days, followed by lysis by direct addition of RIPA buffer. Protein expression was assayed by western blot by using antibodies against TDP-43 (Proteintech, #10782-2-AP, 1:10,000 dilution), HA tag (transgene, GenScript, #A01244, 1:10,000 dilution), and histone H3 (loading control, Abcam, #10799, 1:10,000 dilution).

For subcellular fractionation, transgene expression was induced with 1 µg/ml doxycycline for 7 days, then fractionated according to [67]. 100ng of isolated RNA from each fraction was prepared for sequencing using the KAPA mRNA HyperPrep Kit (Roche, #08098123702) as described above.

## AUTHOR CONTRIBUTIONS

Experiments were conceived by CM, AA, HAR, DD, and JMT and were performed by CM, AA, KFV, BBG, GB, AH, and LBG. Data analysis was performed by CM, AA, KFV, BBG, LBG, DD, and JMT. Funding for the project was secured by HAR, DD, and JMT. The manuscript was written by CM and JMT. The manuscript was edited by CM, DD, HAR, and JMT.

## DATA AVAILABILITY

All high-throughput sequencing data associated with these experiments has been deposited at the Gene Expression Omnibus under accession number GSE288185.

## SUPPLEMENTARY TABLES

**Supplementary table 1.** Differences in RNA localization between wildtype and knockout CAD cells. The log2FC in column J represents log2 fold changes in neurite enrichment (KO - WT). The associated corrected pvalues are found in column L.

**Supplementary table 2.** Differences in RNA localization between wildtype and knockout N2A cells. The log2FC in column J represents log2 fold changes in neurite enrichment (KO - WT). The associated corrected pvalues are found in column L.

**Supplementary table 3.** Count data from MPRA experiment. Read counts are raw read counts. UMI counts are the number of unique UMI counts associated with each oligo.

**Supplementary table 4.** RBNS results. Counts in columns B-I are the raw counts for each oligo in each experiment. Normalized counts in J-Q have been normalized to the total number of counts for that sample.

**Supplementary table 5.** Nucleotide conversion counts for SLAM-seq data. Column C is the number of T to C conversions counted for each oligo, while column D is the number of nonconverted T residues.

**Supplementary file 1.** Fasta file of oligonucleotide sequences used in MPRA, RBNS, and SLAM-seq experiments. Names are of the format <ensembl_gene_id>|<gene_name>|<mutation_location> where mutation locations indicates the positions of the sequence that have been mutated from wildtype, endogenous sequence.

**Supplementary file 2.** Genomic locations (mm10) of oligonucleotide sequences used in MPRA, RBNS, and SLAM-seq experiments.

## SUPPLEMENTARY FIGURES

**Supplementary figure 1.**
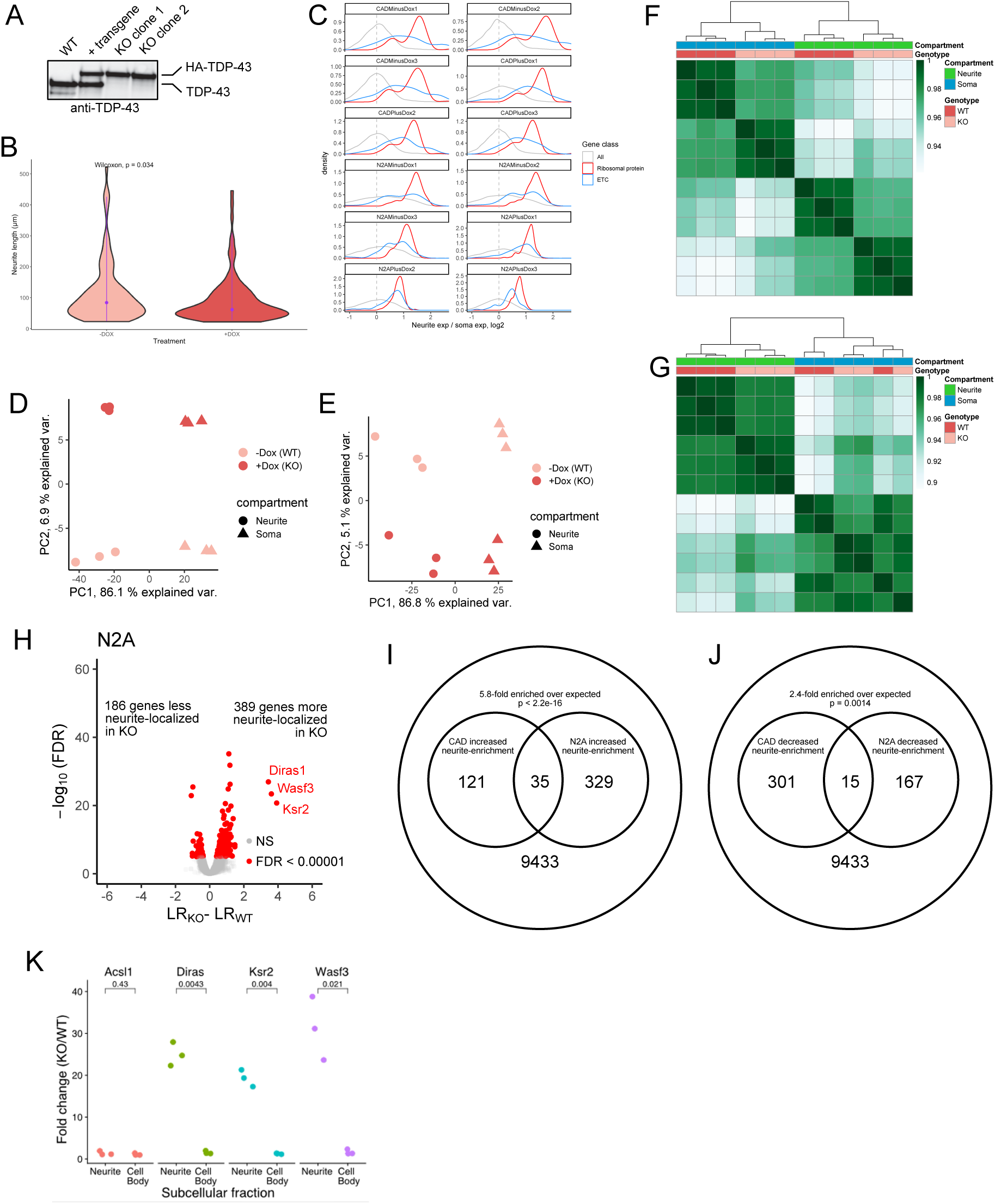
(A) Immunoblot of CAD cells with single copy, site-directed integration of doxycycline-repressible TDP-43. (B) Neurite lengths in CAD cells containing (-dox) or lacking (+dox) TDP-43. (C) Neurite enrichments of RNAs known to be neurite enriched, including those encoding ribosomal proteins and components of the electron transport chain. (D) PCA analysis of RNA expression values from soma and neurite samples of wildtype and TDP-43 knockout CAD cells. (E) As in C, but for N2A cells. (F) Hierarchical clustering of RNA expression values from soma and neurite samples of wildtype and TDP-43 knockout CAD cells. (G) As in E, but for N2A cells. (H) Changes in RNA localization to neurites in N2A cells upon loss of TDP-43. (I) Overlap in the identities of RNAs with increased neurite enrichments upon TDP-43 knockout in CAD and N2A cells. (J) As in G, but for RNas with decreased neurite enrichments. (K) Changes in neurite localization for Ascl1, Diras1, Ksr2, and Wasf3 RNAs between wildtype and knockout CAD cells as measured by RT-qPCR.

**Supplementary figure 2.**
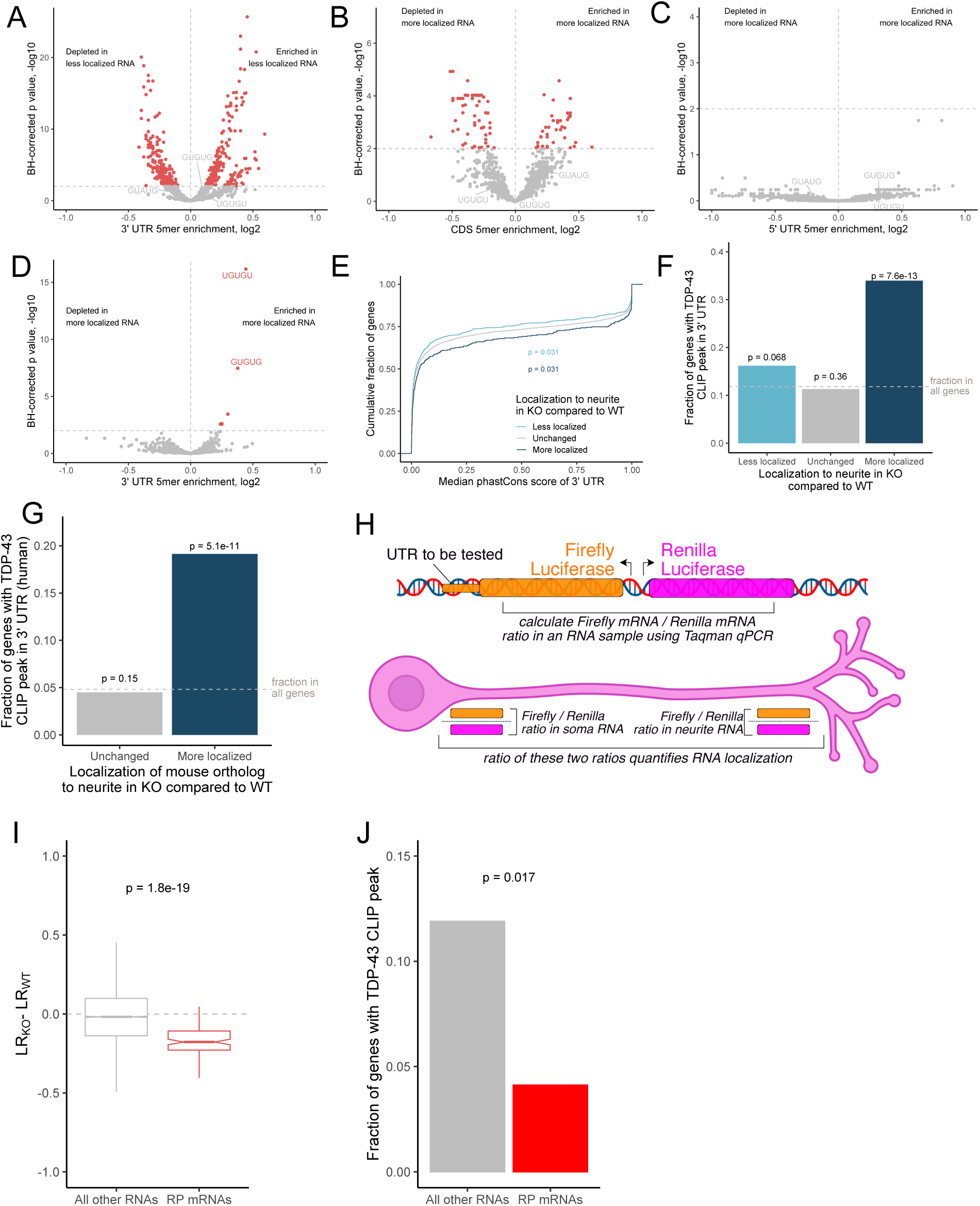
(A) Enrichment for all 5mers in the 3′ UTRs of RNAs that were less neurite localized upon TDP-43 knockout. (B) Enrichment for all 5mers in the coding regions of RNAs that were more neurite localized upon TDP-43 knockout. (C) Enrichment for all 5mers in the 5′ UTRs of RNAs that were more neurite localized upon TDP-43 knockout. (D) Enrichment for all 5mers in the 3′ UTRs of the human orthologs of RNAs that were more neurite localized upon TDP-43 knockout. (E) Phastcons scores for 3′ UTRs of RNAs with the indicated changes in neurite localization upon TDP-43 knockout. (F) Fraction of RNAs with CLIP-seq peaks in their 3′ UTRs for RNAs with the indicated changes in neurite localization upon TDP-43 knockout. (G) As in F, but for the human orthologs of the genes in F. (H) Schematic of reporter RNAs used in experiments monitoring RNA localization of reporters by RT-qPCR. (I) Differences in neurite localization between wildtype and knockout cells for all RNAs (left) or those encoding ribosomal proteins (right). (J) Fraction of genes with a TDP-43 in their 3′ UTR for all RNAs (left) or those encoding ribosomal proteins (right). P values for panels E and I were calculated using a Wilcoxon ranksum test while those in panels F, G, and J, were calculated using a binomial test.

**Supplementary figure 3.**
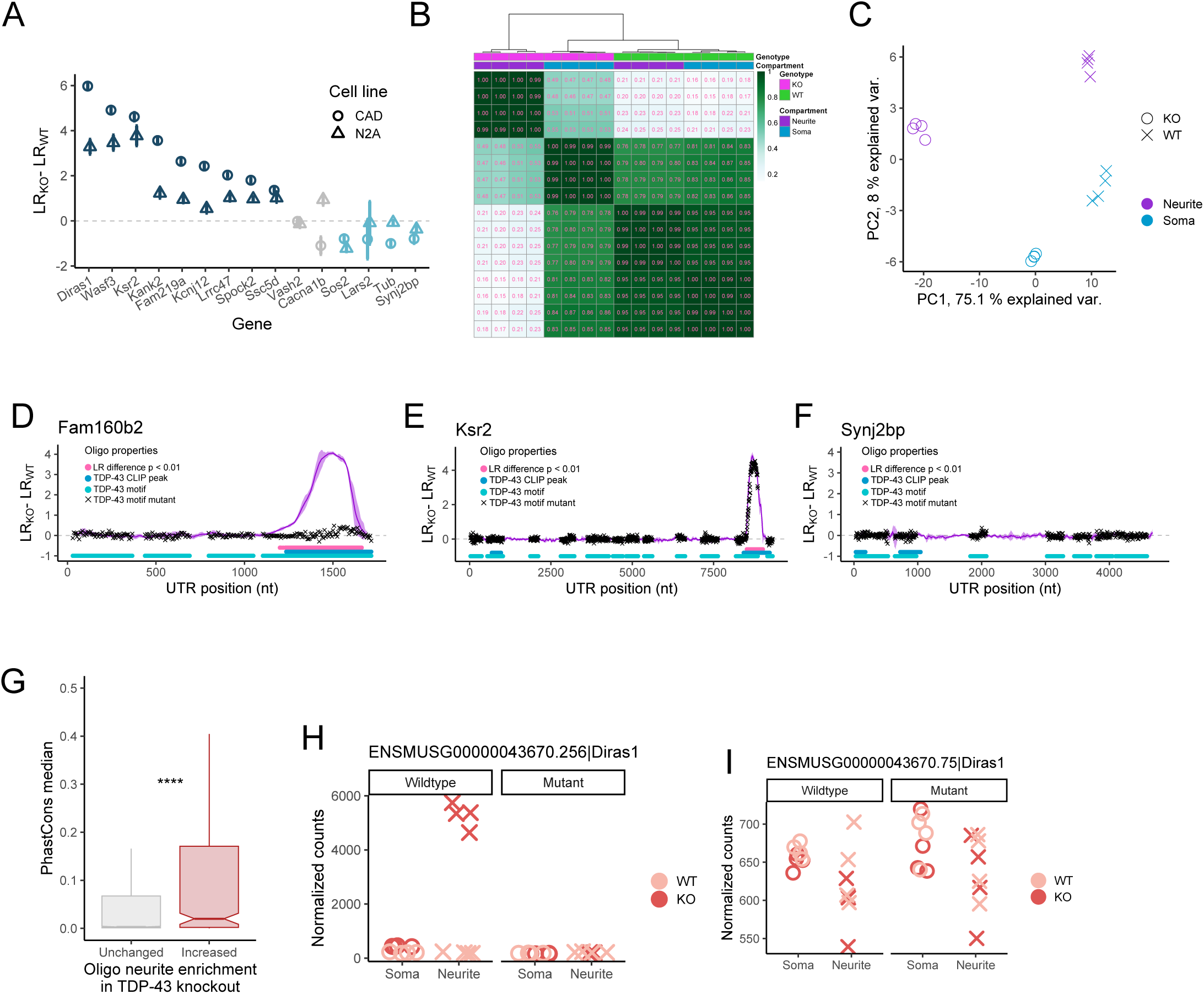
(A) Neurite enrichments for genes chosen for further analysis by MPRA. (B) Hierarchical clustering of normalized expression counts for all MPRA reporters. (C) PCA analysis of normalized expression counts for all MPRA reporters. (D) Differences in neurite enrichment between wildtype and knockout cells for all oligos that tile across the Fam160b2 3′ UTR. Oligos that contain a TDP-43 motif are represented by a light blue circle, those that contain a TDP-43 CLIP-seq peak are represented by a dark blue circle, and oligos with mutated TDP-43 motifs are represented by a black x. (E) As in D, but for the Ksr2 RNA. (F) As in D, but for the non-TDP-43-target Synj2bp RNA. (G) Conservation levels for oligo sequences that were not sufficient to drive TDP-43-dependent changes in RNA localization (left) and those that were sufficient (right). (H) Example of one oligo that contained a TDP-43 motif and lied within a TDP-43 CLIP-seq peak. (I) Example of one oligo that contained a TDP-43 motif but did not lie within a TDP-43 CLIP-seq peak. P values were calculated using Wilcoxon ranksum tests. NS (not significant) represents p >0.05, * p < 0.05, ** p < 0.01, *** p < 0.001, and **** represents p < 0.0001.

**Supplementary figure 4.**
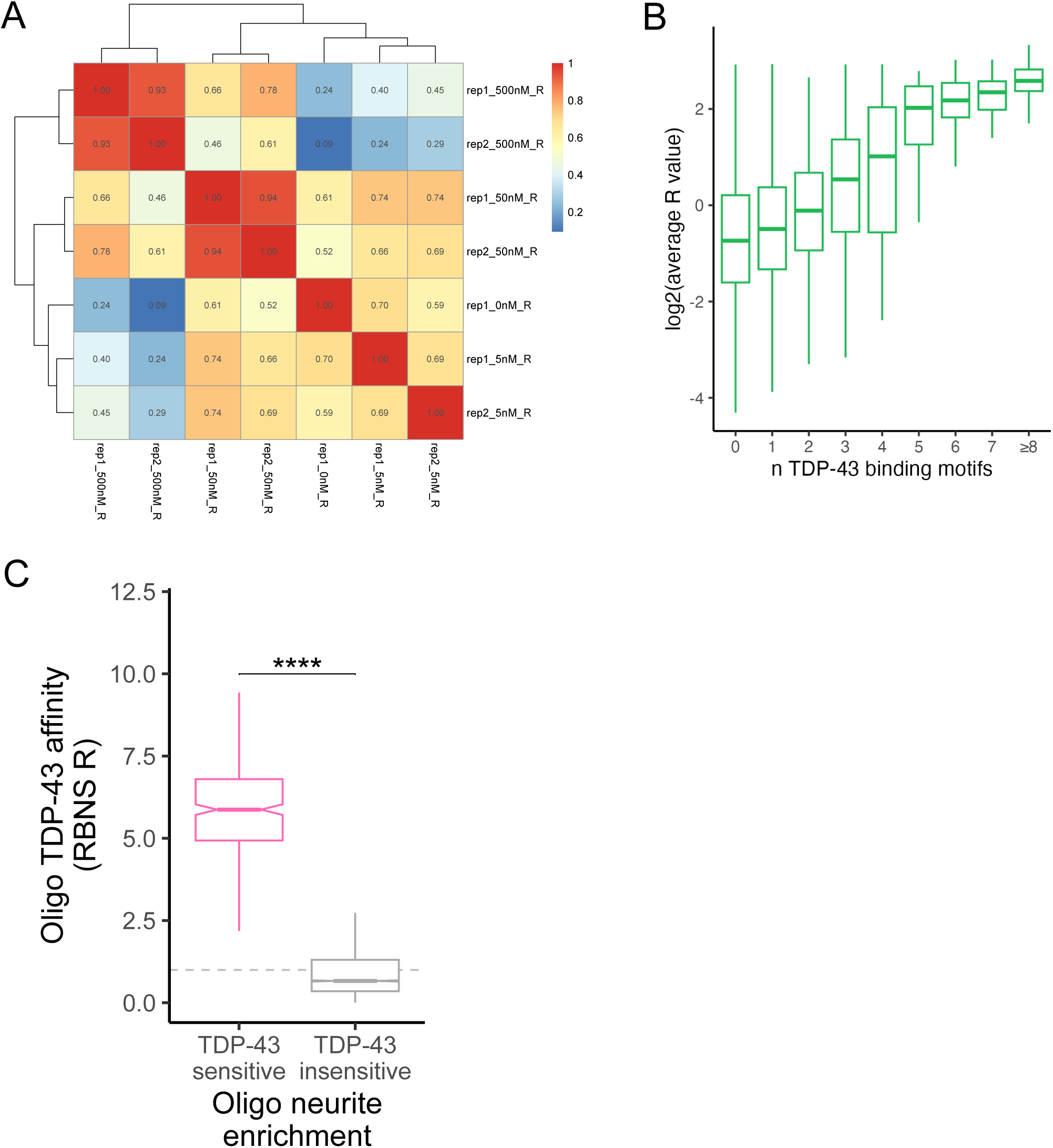
(A) Hierarchical clustering of TDP-43 affinity values for all 260mer sequences as defined by RBNS. (B) Number of TDP-43 motifs in each 260mer sequence and the affinity of that sequence for TDP-43. (C) Differences in TDP-43 affinity for RNA sequences that were sufficient to drive TDP-43-dependent changes in RNA localization (left) and those that were not (right). P values were calculated using Wilcoxon ranksum tests. NS (not significant) represents p >0.05, * p < 0.05, ** p < 0.01, *** p < 0.001, and **** represents p < 0.0001.

**Supplementary figure 5.**
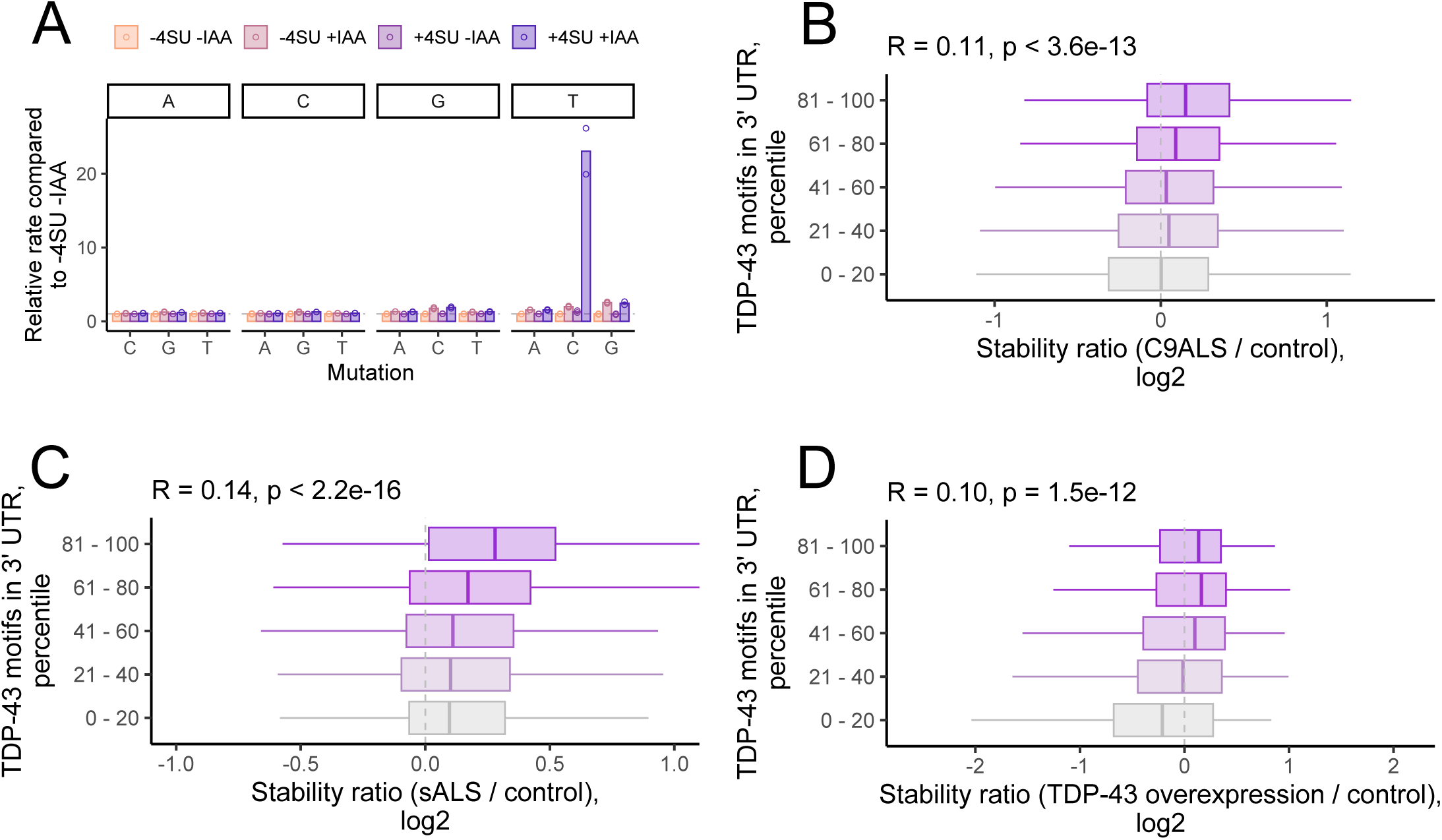
(A) Nucleotide conversion rates for SLAM-seq samples containing or lacking 4SU and iodoacetamide (IAA). (B) Changes in stability as identified by Bru-Chase-Seq between C9ALS and control samples for RNAs with the indicated TDP-43 motif content in their 3′ UTRs. (C) As in B, but for changes in stability between sALS and control samples. (D) As in B, but for changes in stability between TDP-43 overexpression and control samples. The indicated correlation coefficients are Spearman coefficients.

**Supplementary figure 6.**
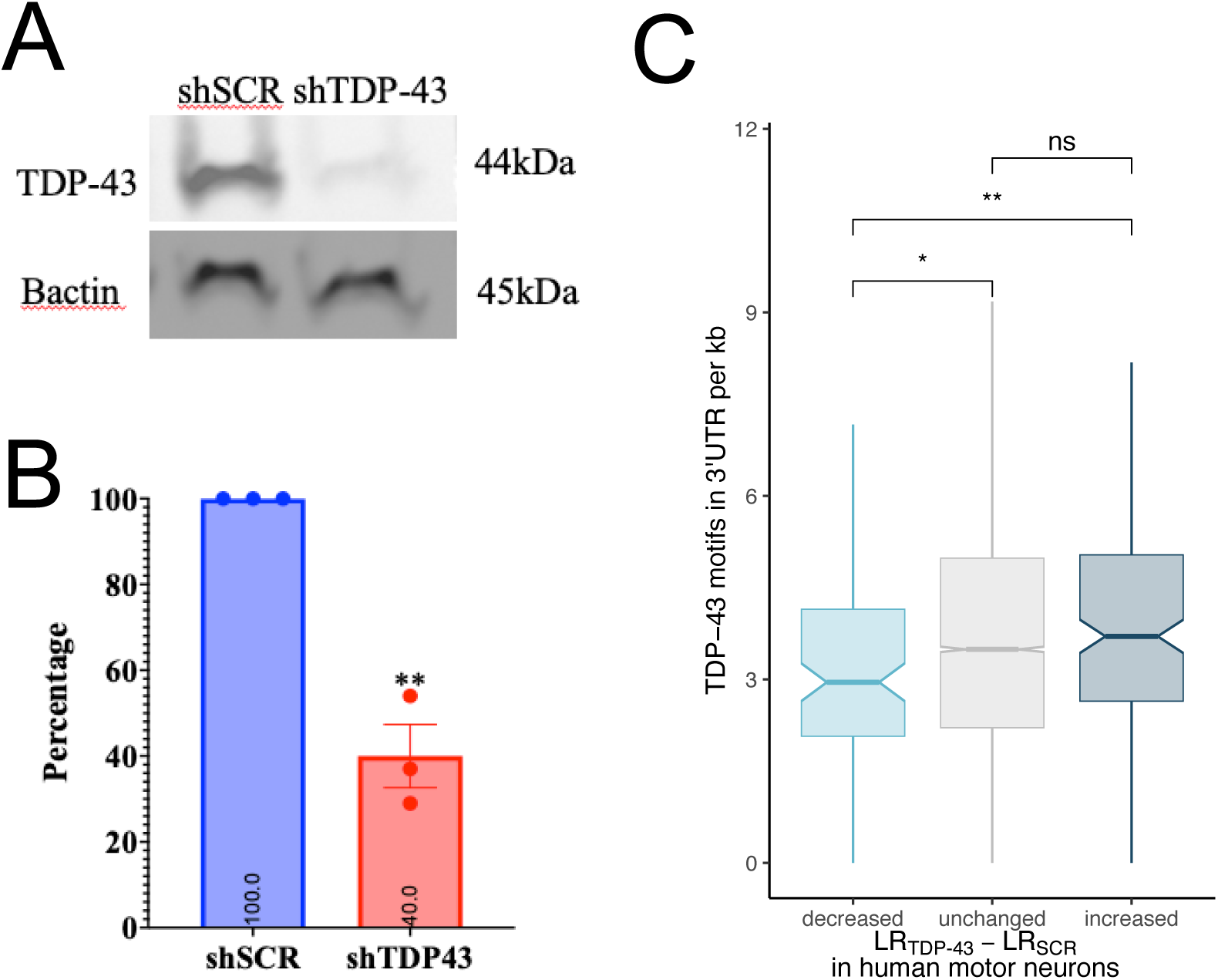
(A) Immunoblot of TDP-43 in human motor neuron samples treated with shRNA against TDP-43 or a control scrambled RNA. (B) Quantification of TDP-43 levels in TDP-43 knockdown experiments. (C) TDP-43 motif content in the 3′ UTRs of RNAs with the indicated changes in neurite localization in human motor neurons between samples treated with TDP-43 siRNA and those treated with control, scrambled siRNA.

